# The usage of transcriptomics datasets as sources of Real-World Data for clinical trialling

**DOI:** 10.1101/2022.11.10.515995

**Authors:** Pedro Matos-Filipe, Juan Manuel García-Illarramendi, Guillem Jorba, Baldo Oliva, Judith Farrés, José Manuel Mas

## Abstract

**Background:** Randomised Clinical Trials (RCT) reflect results within their specific controlled settings, necessitating further studies to understand outcomes across all possible scenarios. The usage of Real-World Data (RWD) has been recently considered to be a viable alternative to overcome these issues and complement clinical conclusions. Molecular profiles of patients captured by high-throughput measures reflect their medical conditions. When this information is linked to clinical and demographical information, nuances in transcriptomics data can uncover subtle variations in disease pathways among distinct patient groups. This work focuses on the construction of a patient repository database with molecular and clinical information resulting from the integration of publicly available transcriptomics datasets.

**Results:** Patient data were integrated into the patient repository by using a novel post-processing technique allowing for the usage of samples originating from different/multiple Gene Expression Omnibus (GEO) datasets. Our post-processing technique, which we have named MicroArray Cross-plAtfoRm pOst-prOcessiNg (MACAROON), aims to standardise and integrate transcriptomics data (considering batch effects and possible processing-originated artefacts). This process was able to better reproduce the down streaming biological conclusions in a 45% improvement compared to other methods available. Furthermore, RWD was mined from GEO samples’ metadata and a clinical and demographical characterisation of the database was obtained. RWD mining was done through a manually curated synonym dictionary allowing for the correct assignment (95.33% median accuracy; 80.14% average) of medical conditions.

**Conclusions:** Our strategy produced a repository, which includes molecular, clinical and demographical RWD by integrating multiple public datasets. The exploration of these data facilitates the discovery of clinical outcomes and molecular pathways specific to predetermined patient populations.

## 1 Introduction

Developments in the fields of Artificial Intelligence (AI) and Machine Learning (ML) (and their association with the pharmaceutical industry)have helped to boost the time-line of production of pharmaceutical products, leading to a reduction in cost (Jayatunga et al., 2022). Techniques in the field are already being streamlined into production under various stages of the life cycle of pharmaceutical products (Paul et al., 2021), namely, drug discovery (Koscielny et al., 2017; Jiménez-Luna et al., 2020; Rodriguez et al., 2021), pharmaceutical development (Rashid et al., 2018), and clinical trial development and monitoring (Kolla et al., 2021; Collins et al., 2021). Nonetheless, regulatory agencies still mostly admit evidence acquired through Randomised Clinical Trials (RCT). At the same time, these institutions encourage the use of stored information about patients and molecular processes to reduce the number of RCTs whenever possible. The combination of molecular data linked to clinical phenotypes can render meaningful patterns. When molecular data are available, it is crucial to as certain the relationship between the known clinical phenotypes and the molecular profile of patients. Extracting meaningful clinical patterns from molecular information can be beneficial. However, obtaining molecular data in RCTs may present practical challenges. These challenges may arise due to difficulties in recruiting patients, particularly for rare diseases. Additionally, RCTs may not always reflect observable results out of controlled settings, necessitating further lengthy observational studies (Andre et al., 2020). The usage of Real-World Data (RWD) has been recently considered to be a viable alternative to complement clinical conclusions. Despite some divergences (i.e., specific data sources), RWD is defined as data relating to patients’ health status that did not originate in traditional clinical trials. Since 2018, both the US Food and Drug Administration (FDA) (Food and Administration, 2018) and the European Medicines Agency (EMA) (Cave et al., 2019) have laid down their regulatory frameworks from which the usage of RWD could be used to obtain valid clinical evidence. In both cases, the accuracy and reliability of RWD (defined according to the scope of the study (Food and Administration)), as well as consistency in data handling and analysis methods (Thorlund et al., 2020), were clear requirements for its acceptance for regulatory purposes (Cave et al., 2019; Zura et al., 2021).

In 1970, Francis Crick presented the Central Dogma of Biology (Crick, 1970).According to his theory, information that codes proteins in living organisms flows from the genome, establishing the relationship between the individuals’ genome and the phenotype it would display. Although the current paradigm now includes a more dynamic approach in comparison to the one Crick presented, the main idea that genetic material influences the proteome and, consequently, the remaining molecules contained within the organism, remains unrefuted (Shapiro, 2009). Therefore, the analysis of an organism’s genome and transcriptome sheds light on molecular events underlying human biology and disease (Melé et al., 2015), producing significant insight into its phenotype, and giving key clues into how to influence it.

The advent of High Throughput (HT) biological techniques in the 1990’s made available the analysis of multiple targets at once, from multiple perspectives (e.g., proteomics, metabolomics, genomics). HT techniques like Microarrays (MA) (1997) (Velculescu et al., 1997) and RNA Sequencing (RNA-Seq) (2006) (Bainbridge et al., 2006) allow the characterisation of the samples’ transcriptome in a quick way that is light on labour and costs. When compared to MA, RNA-Seq presents some advantages concerning sensitivity (Fu et al., 2009) and dynamic range^1^ (Zhao et al., 2014) and is particularly suited for *de novo* sequencing since it does not require reference transcripts for probe design (Wang et al., 2009). On the other hand, MA shows higher throughput, is less labour intensive (Minnier et al., 2018) and, due to being an older technology, presents more currently available data (Thompson et al., 2016) which is a key aspect when dealing with big data studies (Belle et al., 2015; Roski et al., 2014). Gene expression values seem to correlate in both techniques (Zhao et al., 2014; Chen et al., 2017; Nookaew et al., 2012), which validates the usage of any of them if limitations are considered.

The usage of AI, and specifically ML algorithms, is dependent on high volumes of data (Park et al., 2021). The great volumes of data available today led to the adoption of a culture of sharing and reusing HT-generated information within the scientific community, and to the development of public biological databases to store them. After 30 years, the amount of data hosted yearly in these repositories reaches Terabytes, and it is expected to reach Exabytes by 2025 – surpassing data created by social media platforms (Stephens et al., 2015). Several scientific works were published exploring the usage of transcriptomics datasets as RWD sources (Arisi et al., 2019; Kato et al., 2020; Fernandes et al., 2021). Table 1 lists four databases containing transcriptomics experimental data. The Gene Expression Omnibus (GEO) (Barrett et al., 2013) is, to our knowledge, the database with the highest number of available samples and experiments. However, a high number of entries on the platform only include data already pre-processed by the submitting author. A large variety of methods to background correct, normalise and summarise transcriptomics data (Chen and Zhou, 2016) are being used, with no consensus. To avoid this issue, most current protocols for data integration involve the usage of raw data (Carey et al., 2018; Junet et al., 2021) and samples from the same experiment (Koeppen et al., 2017). As last resort, the usage of samples involving the same techniques, machines and processing methods is also commonly used (e.g., GEO datasets (Barrett et al., 2013)). Consequently, a large portion of data included in these transcriptomics repositories is usually excluded from data integration methodologies and protocols. Furthermore, the metadata accompanying the processed files often consists of inaccurate information regarding the processing procedures which completely invalidates its usage; or contains too complex vocabulary (Chen et al., 2019; Khomtchouk et al., 2018) grammar and sentences, hindering its usage in experiments relying on BigData given their lack of machine interpretability (Hadley et al., 2017). Nonetheless, if properly processed and transformed, biological datasets mined from GEO can be used in its fullest and serve as RWD sources. By exploring the association between molecular information (harvested from gene expression values) and clinical variables (obtained from metadata reported in each sample), it is possible to obtain accurate evidence to support a clinical outcome upon a given treatment.

**Table 1.** Biggest genome databases and the number of available human samples and experiments (data obtained from each platform on the 3rd of June 2021). * ”NA” indicates that the data was not available on the platform.

| Database |  | Number of Samples | Number of Experiments | Ref |
| --- | --- | --- | --- | --- |
| *Gene Expression Omnibus (GEO)* |  | 2,293,223 | 66,569 | (Barrett et al., 2013) |
| *ArrayExpress* |  | 1,309,771 | 27,068 | (Brazma et al., 2003) |
| *Japanese Genotype-phenotype Archive (JGA)* |  | 278,601 | NA | (Fukuda et al., 2021) |
| *The European Genome-phenome Archive (EGA)* |  | NA | 4,432 | (Lappalainen et al., 2015) |

This paper presents a novel method for analysing GEO MA studies to obtain clinical data. Given the variability in HT data, we developed a technique called MicroArray Cross-plAtfoRm pOst-prOcessiNg (MACAROON) to correct and standardise MA data from different experiments and processing methods. First, we used text mining to standardise clinical labels in the HT repository and identify only patient data, excluding non-human and immortalised cell line data. This information is later analysed alongside language processing techniques to build ML models that clarify original data transformations and address poor metadata issues. Next, we propose a protocol that uses HK genes’ expression values to adjust for different normalisation techniques applied to MA datasets before they were uploaded to GEO. This approach allows researchers to combine MA profiles from multiple sources, regardless of the samples’ pre-processing steps. Additionally, text mining standardises metadata by classifying samples based on health status and demographics, creating a source of RWD. This method allows to link this RWD with molecular patterns by exploring the transcriptomic data from HT experiments. With this effort, we aim at easing access to RWD, which is currently a main limitation of pharmaceutical companies in this kind of studies (Wallach et al., 2021).

## 2 Results and Discussion

The foremost objective of this work is to exploit the GEO database (Barrett et al., 2013) as a source of molecular and phenotypical data (RWD). We have obtained a total of 1,866,292 MA samples (including gene expression and corresponding metadata). Gene expression data were subjected to the novel post-processing method MACAROON (introduced in this work), while clinical metadata was mined for phenotypical labels (demographics and medical conditions).

Clinical labelling in GEO is often restricted to medical conditions of interest to the specific experiment to which they belong. Consequently, further medical conditions may be disregarded, even in cases in which these are highly probable. For instance, patients with ageing-associated conditions usually display other disorders of the same type. As such, phenotypical labels in combination with the corresponding gene expression data were used to generate classifiers aimed at predicting clinical phenotypes excluded from metadata. The whole set of samples was run through these classifiers, and, for each sample, newly predicted clinical labels were generated – further complementing our RWD repository.

In summary, each sample in the RWD repository consists of the post-processed MA data, and the corresponding clinical and demographical labels. To better guide the reader through the methods presented herein, an overall depiction of the pipeline deployed in this work is schematized in figure 1. This new resource links the genotype of the patient (collected from the molecular description included in the transcriptomics data – figure 1A) to its phenotype (gathered from the clinical and demographical data – figure 1B). Phenotypical data includes information mined from GEO files (figure 1B.1) and complemented with ML predictions of further clinical labels (figure 1B.2).

**Fig. 1.**
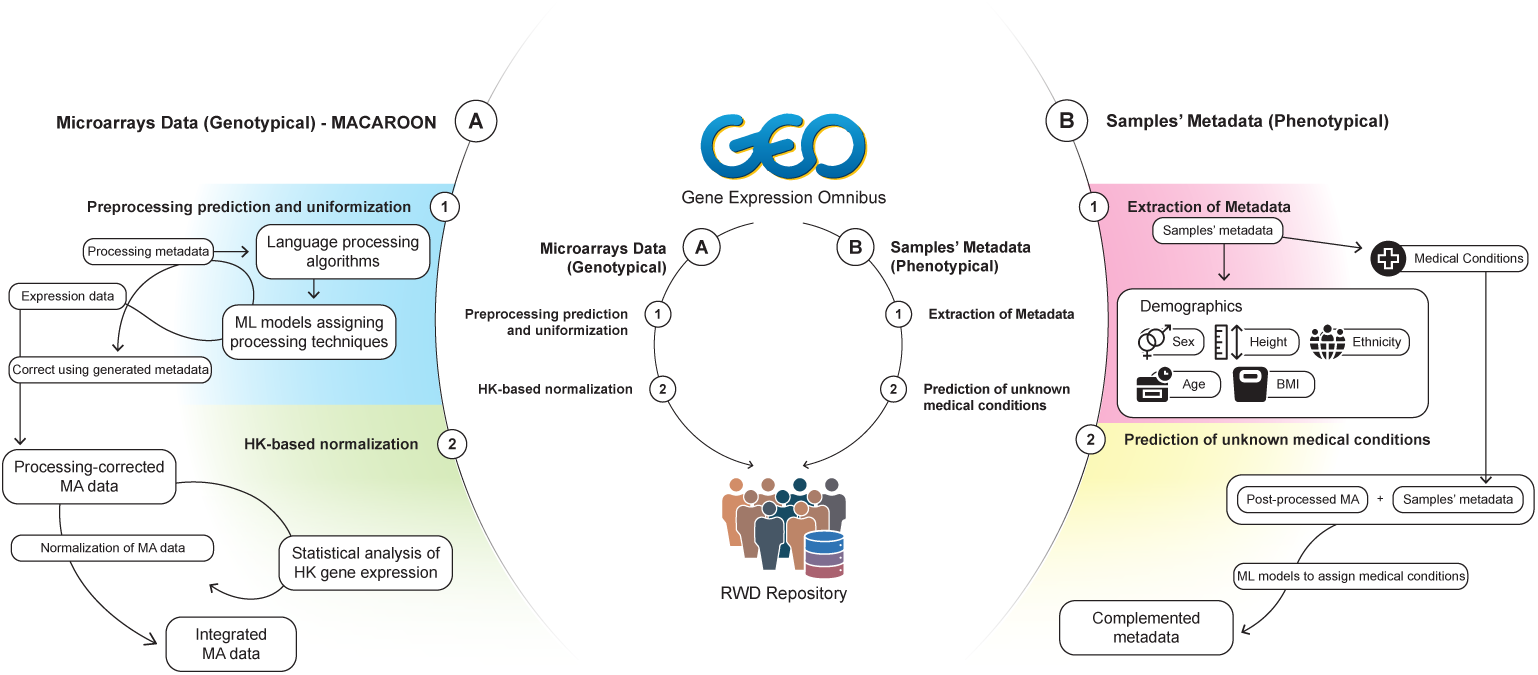
The overall pipeline of the methods applied in this work. (A) Collection and processing of MA data. (B) Collection and processing of phenotypical data from GEO.

### Global microarray data processing

MACAROON is split into two stages. First, we revert and clean the processing procedures applied by the original researcher (i.e., reversal of logarithm procedures and labelling of non-intensity files) to uniformise the orders of magnitude in the samples’ data. Then, we apply a GEO-wide normalisation process to reduce the impact of sample-specific background corrections – which could lead to misleading biological conclusions.

### ML-based assignation of samples’ pre-processing

Metadata accompanying the MA samples were downloaded for all GEO samples. Each text file was analysed to label samples for the presence of terms related to data processing, like the application of different logarithms, common MA normalisation methodologies and types of data (i.e., intensity or Differential Expression (DE) results) – that we will describe as “pre-processing” in the rest of this paper. A combination of interest terms was used to select samples in datasets to train ML classifier models (combined into ensemble models) – that we will call “ML_processing_” moving forward. These terms were picked to maximise the distinguishability of “applicable” from “non-applicable” classes (when a processing method was applied or when that specific method was not applied, respectively) – see Methods section for details (table 4). This process resulted in five different MA datasets, each with 5000 samples (70-30 train/test split): application of (1) Log2, (2) Log10, (3) Log (unknown base), (4) DE ratios and (5) normalisation. We then used these datasets for training ML_processing_ for inferring which pre-processing methods were applied to MA samples whose metadata was inconsistent or non-existent. From each sample in the dataset, we extracted a statistical description (minimum, maximum, mean, standard deviation, and variance) and the values of 5 low-variability HK genes (top five genes in table 3 in the Methods section).

**Table 3:** List of HK genes included on the datasets used in this work organised by standard deviation of expression across multiple tissues. The top 5 genes of this list were used in the datasets for sample pre-processing classification.

| Uniprot | Gene Name | Protein Function | $\sigma(\log_2((RPKM)))$ |
| --- | --- | --- | --- |
| *Q99497* | PARK7 | Parkinson’s disease protein<br>7 | 0.273137 |
| *Q15843* | NEDD8 | NEDD8 | 0.350249 |
| *P28070* | PSMB4 | Proteasome subunit beta<br>type-4 | 0.370264 |
| *P84090* | ERH | Enhancer of rudimentary<br>homolog | 0.382595 |
| *P28074* | PSMB5 | Proteasome subunit beta<br>type-5 | 0.397559 |
| *P19338* | NCL | Nucleolin | 0.423427 |
| *P78344* | EIF4G2 | Eukaryotic translation initi-<br>ation factor 4 gamma 2 | 0.446228 |
| *P84077* | ARF1 | ADP-ribosylation factor 1 | 0.458529 |
| *P15954* | COX7C | Cytochrome c oxidase sub-<br>unit 7C, mitochondrial | 0.474006 |
| *P49773* | HINT1 | Histidine triad nucleotide-binding protein 1 | 0.474979 |

**Table 3: List of HK genes included on the datasets used in this work organised by standard deviation of expression across multiple tissues. (*continued*)**
| Uniprot | Gene Name | Protein Function | $\sigma(\log_2((RPKM)))$ |
| --- | --- | --- | --- |
| *P50395* | GDI2 | Rab GDP dissociation inhibitor beta | 0.496972 |
| *P00441* | SOD1 | Superoxide dismutase [Cu-Zn] | 0.502851 |
| *P37108* | SRP14 | Signal recognition particle 14 kDa protein | 0.504101 |
| *P61803* | DAD1 | Dolichyl-diphosphooligosaccharide-protein glycosyltransferase subunit DAD1 | 0.519353 |
| *P18583* | SON | Protein SON | 0.536725 |
| *P0CG47* | UBB | Polyubiquitin-B | 0.537386 |
| *P00558* | PGK1 | Phosphoglycerate kinase 1 | 0.547633 |
| *Q06830* | PRDX1 | Peroxiredoxin-1 | 0.549351 |
| *P48556* | PSMD8 | 26S proteasome non-ATPase regulatory subunit 8 | 0.552184 |
| *Q92572* | AP3S1 | AP-3 complex subunit sigma-1 | 0.581663 |
| *P09429* | HMGB1 | High mobility group protein B1 | 0.595868 |

**Table 3: List of HK genes included on the datasets used in this work organised by standard deviation of expression across multiple tissues. (*continued*)**
| Uniprot | Gene Name | Protein Function | $\sigma(\log_2((RPKM)))$ |
| --- | --- | --- | --- |
| *P25705* | ATP5F1 | ATP synthase subunit<br>alpha, mitochondrial | 0.60272 |
| *P48201* | ATP5A1 | ATP synthase F(0) complex<br>subunit C3, mitochondrial | 0.608176 |
| *Q15046* | KARS | Lysine-tRNA ligase | 0.608522 |
| *P00338* | LDHA | L-lactate dehydrogenase A<br>chain | 0.615549 |
| *P36873* | PPP1CC | Serine/threonine-protein<br>phosphatase PP1-gamma<br>catalytic subunit | 0.620001 |
| *Q15366* | PCBP2 | Poly(rC)-binding protein 2 | 0.622055 |
| *P49770* | EIF2B2 | Translation initiation factor<br>eIF-2B subunit beta | 0.625432 |
| *P04844* | RPN2 | Dolichyl-diphosphooligosaccharide-protein glycosyltransferase<br>subunit 2 | 0.637911 |
| *P0DP23* | CALM1 | Calmodulin-1 | 0.648649 |
| *P14406* | COX7A2 | Cytochrome c oxidase sub-<br>unit 7A2, mitochondrial | 0.656231 |
| *P14854* | COX6B1 | Cytochrome c oxidase sub-<br>unit 6B1 | 0.657215 |

**Table 3: List of HK genes included on the datasets used in this work organised by standard deviation of expression across multiple tissues. (*continued*)**
| Uniprot | Gene Name | Protein Function | $\sigma(\log_2((RPKM)))$ |
| --- | --- | --- | --- |
| *P62875* | POLR2L | DNA-directed RNA polymerases I, II, and III subunit RPABC5 | 0.675001 |
| *P0CG48* | UBC | Polyubiquitin-C | 0.677235 |
| *P10644* | PRKAR1A | cAMP-dependent protein kinase type I-alpha regulatory subunit | 0.680635 |
| *O15144* | ARPC2 | Actin-related protein 2/3 complex subunit 2 | 0.693822 |
| *P17844* | DDX5 | Probable ATP-dependent RNA helicase DDX5 | 0.73098 |
| *O95467* | GNAS | Neuroendocrine secretory protein 55 | 0.731569 |
| *P63092* | GNAS | Guanine nucleotide-binding protein G(s) subunit alpha isoforms short | 0.731569 |
| *P84996* | GNAS | Protein ALEX | 0.731569 |

**Table 4.** Composition of datasets after text mining procedures. The “+” on the right illustrated the “applicable” class, and the “-” the “non-applicable” class. The binary matrix indicates if the samples were included depending on the “hot word”. Cells with “1” indicate that samples with the word were included, cells with “0” indicate that the samples were excluded, and cells with “-” indicate that the “hot word” was ignored. * The term “Normalized” is an umbrella term that additionally includes the words “normali”, “mas”, “quantile”, and “rma”.

| Dataset | Log2 | Log10 | Log | Ratio | Raw | Normalized* |
| --- | --- | --- | --- | --- | --- | --- |
| *Log2+* | 1 | 0 | - | - | - | - |
| *Log2-* | 0 | 1 | - | - | - | - |
| *Log10+* | 0 | 1 | - | - | - | - |
| *Log10-* | 1 | 0 | - | - | - | - |
| *Ratio+* | - | - | - | 1 | 0 | - |
| *Ratio-* | - | - | - | 0 | 1 | - |
| *Log+* | - | - | 1 | - | 0 | - |
| *Log-* | - | - | 0 | - | 1 | 0 |
| *Normalized+* | - | - | - | - | 0 | 1 |
| *Normalized-* | - | - | - | - | 1 | 0 |

The datasets mentioned above were used to train ML_processing_ to infer if each determined processing technique was applied to the sample or not. Given the large number of missing values in the training datasets (especially concerning HK MA intensities), the various models were trained with a different set of features, as a result of an automatic feature selection strategy. Given that training datasets were built with class balancing in mind, we analysed models’ results based on accuracy (ACC), corresponding to the proportion of correct predictions provided by the models (Equation 1). In our results, models could attain a satisfactory level of accuracy (25-k fold Cross-validation (CV); in the range [70%-98%]). To maintain a maximum certainty in all predictions, we explored the accuracy of the top-three models in each 5%-quantile of the score range in that model. We found that Log2, Log10, Log and Ratio were capable of outputting statistically significant classifications (95% correct answers per score quantile) in both classes (see Supplementary figure 1). Yet, in Normalisation, ML models did not display satisfactory prediction capabilities due to difficulties in clearly selecting samples on the negative class, which made us disregard them in further steps. Final models for each processing procedure being evaluated comprise an ensemble of the top-three best performing ML models in each category (which allowed us to avoid missing value imputation approaches). Exceptions reside in models predicting Log2 and Log10, where ensembles also contain models to distinguish if Log with an unknown base (log*_x_*) was applied (a total of six models). In Table 2, we display the details and 25-k folds CV performance metrics of each model included in the three ensembles. The ML_processing_ were, then, trained sequentially with k-1 parts and tested using the remaining set (Refaeilzadeh et al., 2009). CV metrics consist of the means of the metrics obtained in each training procedure. Here, three further metrics are shown: (1) precision (PRE), corresponding to the proportion of positive results that are true positive results (eq. 2); (2) sensitivity (SNS), corresponding to the proportion of positives that are correctly identified (eq. 3); and (3) specificity (SPC), corresponding to the proportion of negatives that are correctly identified (eq. 4). We assessed the performance of the various ensembles considering the training dataset, the testing dataset (both following the requirements displayed in Table 4), and against all samples in the complete dataset. Metrics corresponding to the train and test sets present very similar results. However, metrics computed using all the samples available present a significant drop across all metrics (average 44.39%) except for SNS (where differences between train/test datasets and all samples average 0,30% across the various ensembles). We believe that this effect relies on the poor definition of negative classes. Given that the text mining process works by identifying specific words present in the metadata data of the various samples, negative classes may be poorly assigned if the data was pre-processed using a technique that was not mentioned in the corresponding metadata. This problem is found frequently in the literature (Koeppen et al., 2017; Chen et al., 2019; Khomtchouk et al., 2018; Hadley et al., 2017). Further test performance metrics are available in Supplementary Table 1 (training metrics can be found in Supplementary Table 2). A comparable assessment of these results can be found in Figure 2.

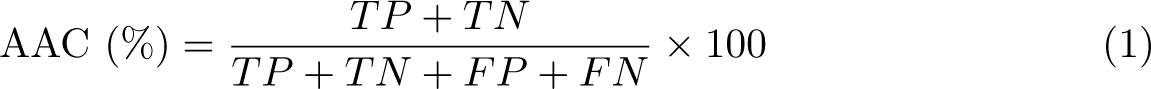

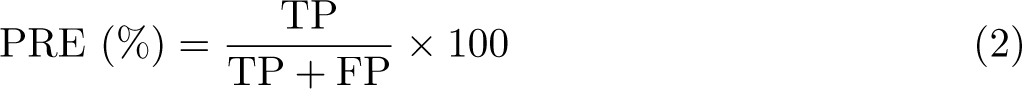

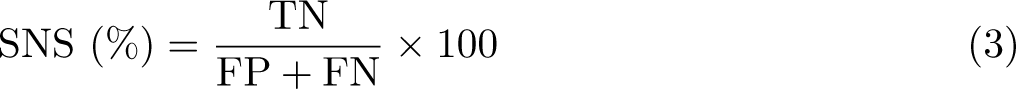

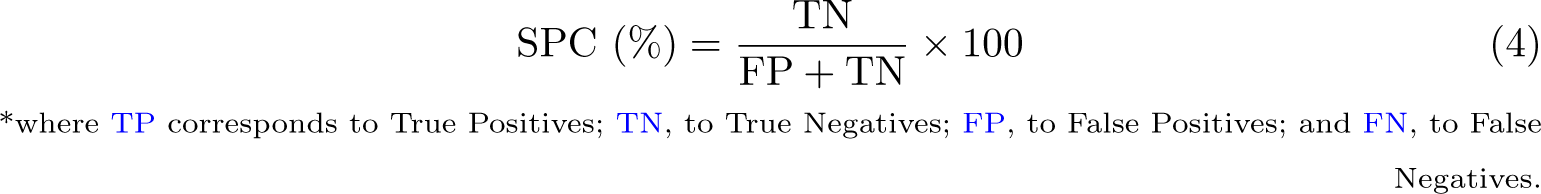

**Fig. 2.**
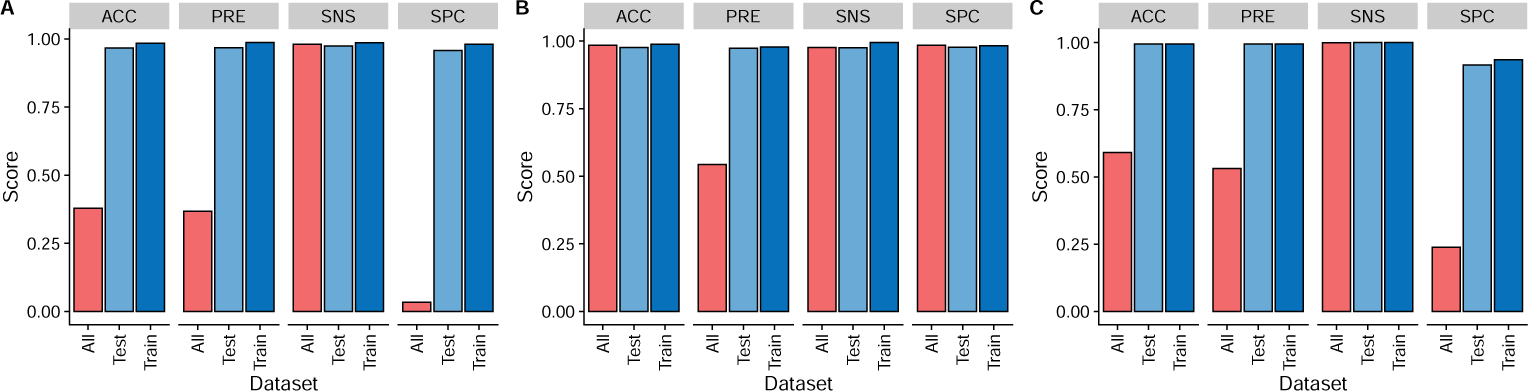
Global test metrics of the various ensemble models in ML_processing_. (A) Log2, (B) Log10, and (C) Ratio.

**Table 2.** Composition of ensemble models for the classification of previous processing.

| Dataset | ML<br>Algo-<br>rithm | Included<br>Features | Feature<br>Selection<br>Method | CV-ACC<br>(%) | CV-PRE<br>(%) | CV-SNS<br>(%) | CV-SPC<br>(%) |
| --- | --- | --- | --- | --- | --- | --- | --- |
| Log2 | SVM | MIN,<br>MAX,<br>AVG,<br>STD,<br>VAR,<br>Q99497,<br>Q15843,<br>P28070,<br>P84090,<br>P28074 | Elastic Net | 94.31 | 91.40 | 96.86 | 92.10 |
|  | SVM | MIN,<br>MAX,<br>AVG,<br>STD,<br>VAR | Simple<br>Regression | 92.99 | 94.24 | 91.92 | 94.11 |
|  | SVM | MIN,<br>MAX,<br>AVG,<br>STD | Random For-<br>est | 92.53 | 93.60 | 91.67 | 93.43 |
| Log10 | SVM | MIN,<br>MAX,<br>AVG,<br>STD,<br>VAR,<br>Q99497,<br>Q15843,<br>P28070,<br>P84090,<br>P28074 | Elastic Net | 94.48 | 96.56 | 92.88 | 96.28 |
|  | SVM | MIN,<br>MAX,<br>STD,<br>VAR | Simple<br>Regression | 92.28 | 91.05 | 93.23 | 91.39 |
|  | SVM | MIN,<br>MAX,<br>AVG,<br>STD,<br>VAR | Wilcoxon | 92.10 | 91.14 | 92.70 | 91.54 |
| Log | TREE | MIN,<br>MAX,<br>AVG,<br>STD,<br>VAR,<br>Q99497,<br>Q15843,<br>P28070,<br>P84090,<br>P28074 | Elastic Net | 98.36 | 97.98 | 98.08 | 98.56 |
|  | TREE | MIN,<br>MAX,<br>AVG,<br>STD,<br>VAR | Ridge | 96.77 | 96.94 | 96.89 | 96.64 |
|  | TREE | MIN,<br>MAX,<br>AVG,<br>STD | Random For-<br>est | 96.47 | 96.54 | 96.73 | 96.19 |
| Ratio | TREE | MIN,<br>MAX,<br>AVG,<br>STD,<br>VAR | Entropy, Cor-<br>relation | 81.13 | 81.00 | 81.48 | 80.78 |
|  | SVM | MIN,<br>STD | None | 78.82 | 71.50 | 96.05 | 61.50 |
|  | SVM | MIN | Elastic Net | 78.39 | 70.70 | 97.20 | 59.49 |

Samples where pre-processing metadata was unclear were classified using the ML_processing_ models. Classification results were used to revert all samples to the original state by reverting the specific Log operation (see Methods section for detail). Given that *Log*(*x*) does not exist if *x* is negative, and to avoid approximation of imaginary numbers, values were restored to their original form. In other words, we have reverted the *Log* transformation in all samples (see 16) instead of uniformising the data by applying *Log*.

**Algorithm 1.**
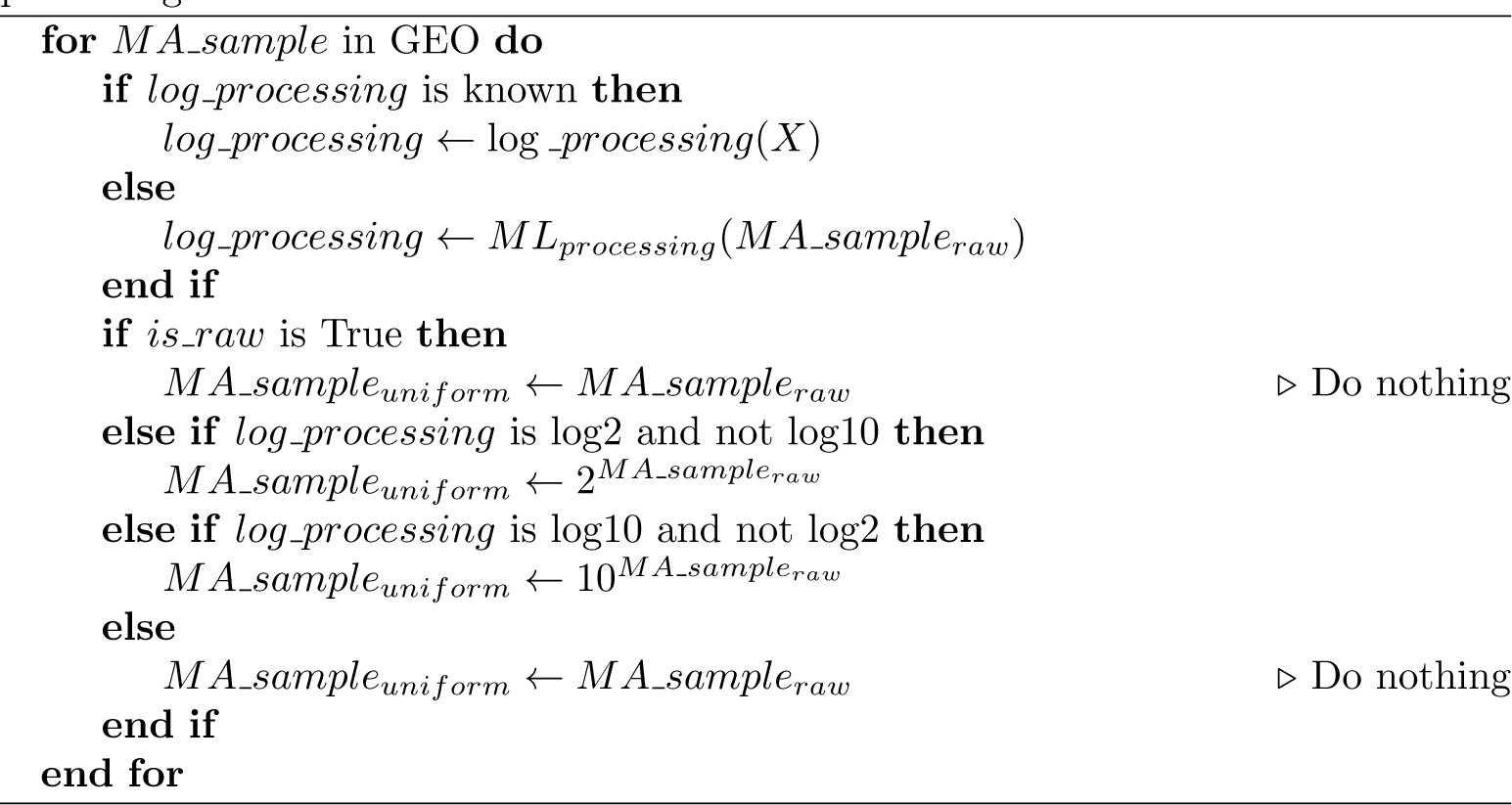
Composition of ensemble models for the classification of previous processing.

### House Keeping-based regression for normalisation

After the first stage of this method – aimed at cleaning and uniformising original processing techniques, we designed a normalisation method based on HK gene expression. The median expression of a set of 40 HK genes selected for their least cross-tissue variability as found in the literature (Eisenberg and Levanon, 2013) was used as reference. For each sample of the whole dataset, genes were mapped in a two-coordinate system and a regression line was fitted using a linear (polynomial regression with one coefficient) and a non-linear (spline regression) method. MA normalisation protocols reliant on regression techniques usually use the LOWESS algorithm (Zahurak et al., 2007). However, this solution could not be used here case as it does not allow for extrapolation. Given that HK genes are rarely under/overexpressed, extrapolation was a requirement.

For benchmarking, a test dataset containing MA data from prostate cancer samples and healthy controls was extracted. This dataset included 394 samples from 63 GEO experiments (as available in Supplementary Table 3). Their expression values were then visually explored according to their distribution sample- and gene-wise; representations of these results are available in figure 3. This figure shows a progressive representation of the datasets as the stages of MACAROON are applied. Taking the general assumption that most genes should not see any change of expression across samples (Vafaee et al., 2019), the mean intensity should show a tendency towards a constant value. This tendency is visible from the raw datasets (figure 3A) to the data after previous pre-processing correction (3B), and then to the data subjected to HK regression (3C, and 3D). The effect is also clear on the MA-plots (bottom of figure 3c), where M is defined by equation 5 and A is defined by equation **??**. With the same premise discussed above, the expression of most genes stays generally constant across samples. Consequently, the M value corresponding to the great majority of genes should stay constant along with A if the data is well normalised. The red line in F3c corresponds to a regression line of the M/A pairs in the plot. If the initial assumption is met, the red and blue lines (corresponding to *M* = 0) should overlap. This is the case for each step of Trans Babel, where the blue and red lines show a tendency to overlap and better depict a biological scenario. On the other hand, samples present a strong localised concentration of data points (see density curves in figures 3A and B). We attribute this event to the lack of dynamic range that is characteristic of MA (Fu et al., 2009).

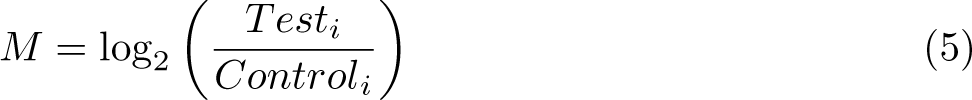

**Fig. 3.**
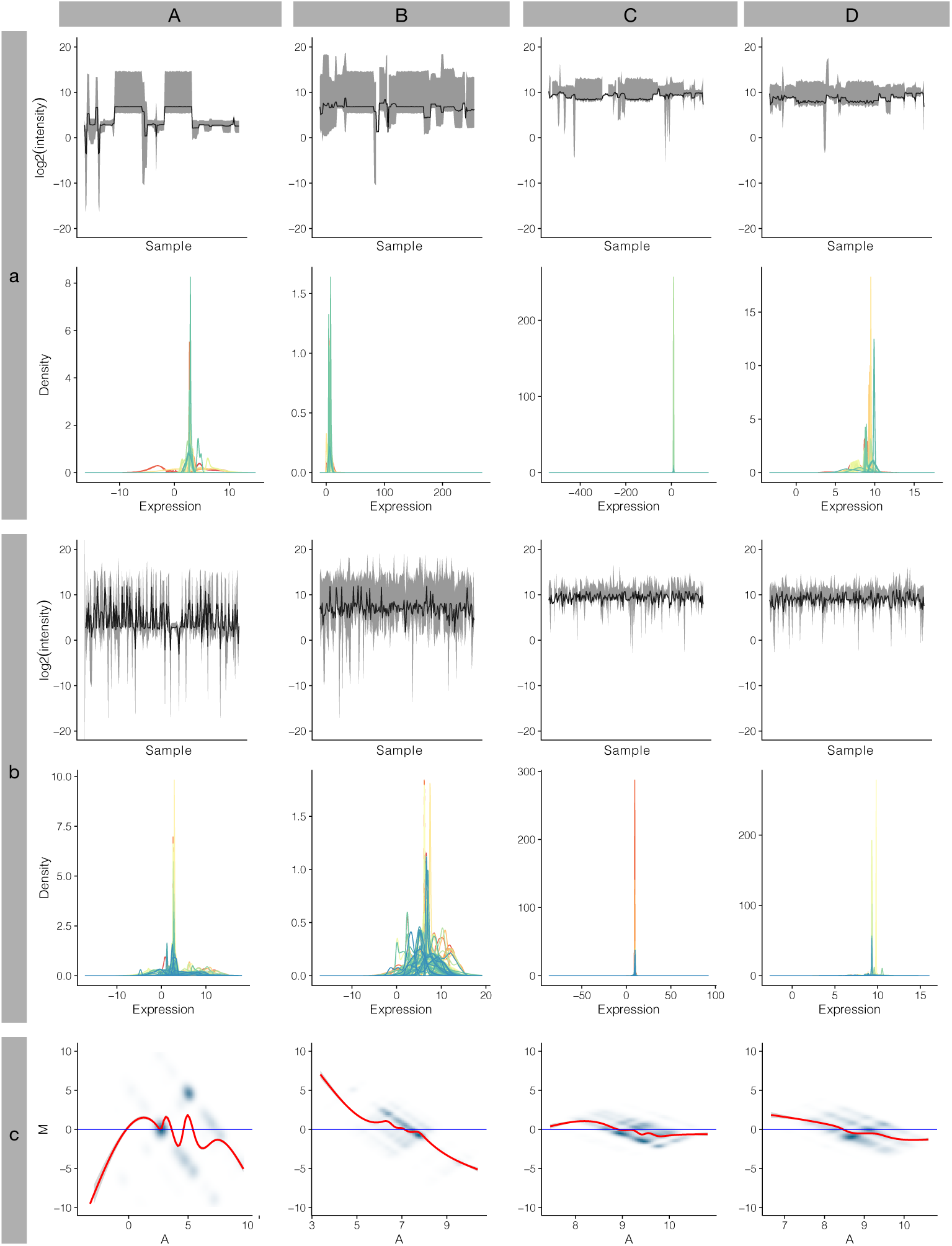
Log-expression data distribution across the various stages of MACAROON. A, B, C and D refer to the datasets produced along the process: A. Raw unprocessed data; B. After reverting preprocessing; C. After spline regression; and D. After polynomial regression. (a), (b), and (c) concern the samples displayed in the figure: (a). healthy control samples; (b). prostate cancer samples; and (c). MA-plot (control vs. prostate cancer) – as described in Eq. 5 and 6.

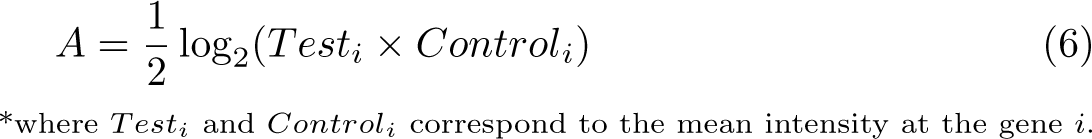

### MACAROON compared against YuGene

As further validation of MACAROON, we compared its output against YuGene (Cao et al., 2014): to our knowledge, the only available method that, like the method presented in this paper, is able to make MA normalisation at the sample level. Figures 4A, B and C present a sequence of bar plots describing the intensity values of the GEO series GSE102124 (Sowalsky et al., 2018) (one of the samples available at the PCa dataset used above in this work) – corresponding to 22 MA samples. The sub-plots correspond to these data in the preprocessed form, followed by the application of MACAROON and YuGene. From the analysis of the figure, we can understand how each one of the methods transforms the raw data and how they impact the final biological interpretability. One of the major drawbacks of MA, in comparison to RNA-Seq, is its low dynamic range. In MACAROON, an effort is made to maintain the standard range of processed MA. On the other hand, in YuGene, values are scaled into a 0-1 range. Given that the output from both methods is equivalent to the *log*_2_ scale, the application of MACAROON provides better equivalence to current methods when performing DE analysis – this fact was later observed when performing a comparative DE analysis with the three datasets. However, MACAROON shows worst performance than YuGene at aligning the samples’ medians, which also impacts the range of DE results. Different MA platforms have different coverage, a concept that portrays the probes that are included and the genes that are assessed by using each platform. Consequently, datasets obtained from the integration of different platforms lead to datasets with many missing values. Our normalisation approach, based on regression, allows the conservation of maximum and minimum values per probe, while supporting missing values, and without imputing them, retaining the original biological motifs in the dataset.

**Fig. 4.**
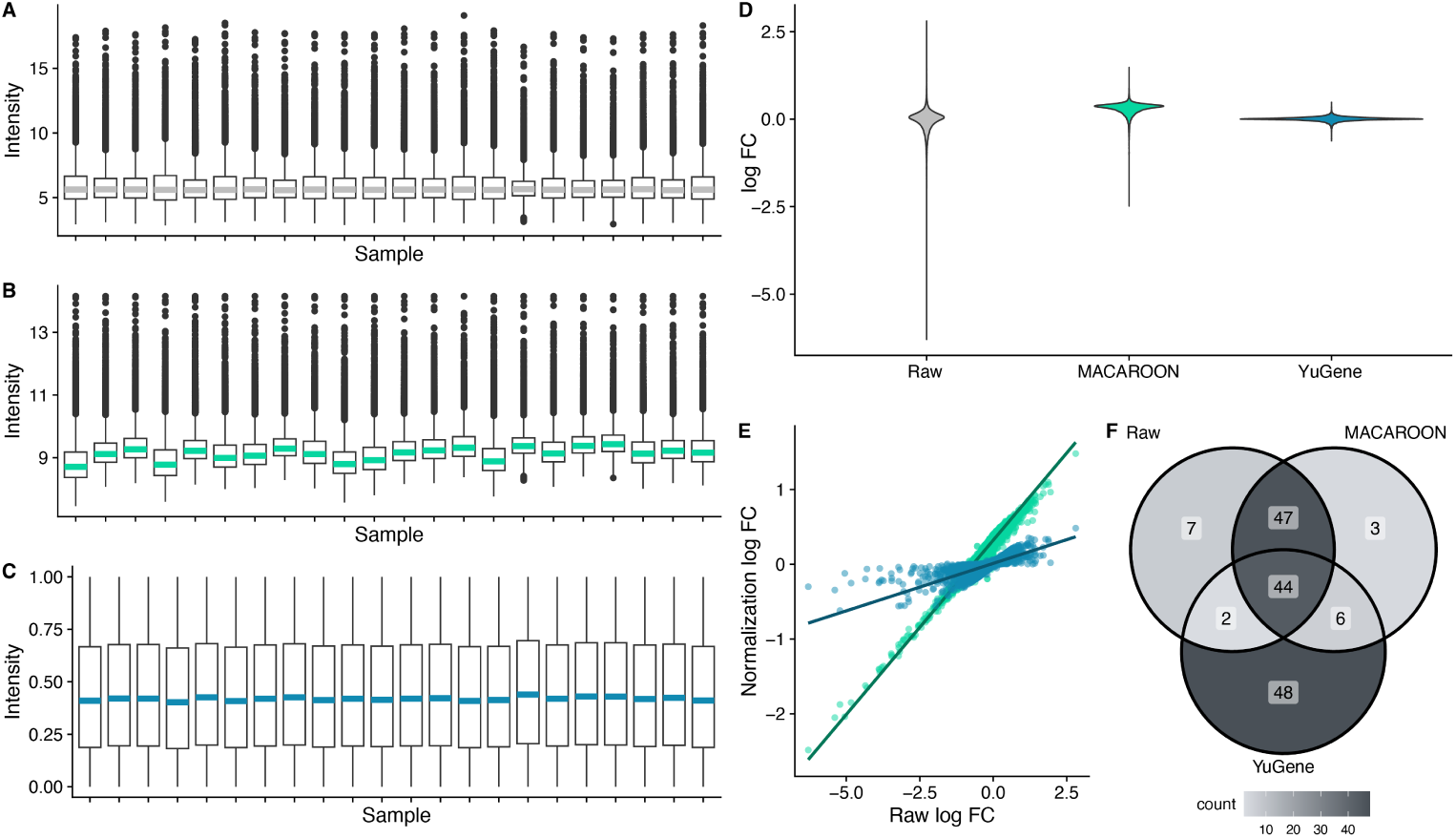
Direct comparison between methods. In the left column, we show the intensity per sample. A corresponds to the *log*_2_ raw data; B, using MACAROON; and C, using YuGene (Cao et al., 2014). In the right column, we display a comparison of the results of a Differential Expression analysis. In D, we show a dispersion of the log Fold Change values; in E, a scatter plot correlating the fold change values per gene using MACAROON and YuGene vs. the values calculated using the *log*_2_ Raw dataset; in F, a Venn diagram portraying common (ttop-50 and bottom-50 DE genes in log FC) differentially expressed genes depending on the method used. When applicable, the elements in grey correspond to the *log*_2_ Raw dataset, in green to MACAROON, and in blue to YuGene.

As means of further comparing the three methods, we ran a DE Analysis with data processed using the different methods (*log*_2_ Raw, MACAROON, YuGene). In figure 4D, we represent the results’ space (density of log Fold-change (FC)) of the DE according to the dataset. As discussed previously, MACAROON and YuGene tend to provide results in a reduced range (in comparison to the original dataset) because of the misalignment of samples’ means or by the heavy scaling of datasets, respectively. Nonetheless, when looking at the correlation between log Fold-change (FC) results (figure 4E), as well as overlapping differentially expressed genes (figure 4F), MACAROON shows greater similarity to the original results (91% matching) in comparison to YuGene (46%), suggesting better biological interpretability.

### Case Study: Differential Analysis Experiment

The integrated PCa dataset presented earlier (available in Supplementary Table 3) was used again to present the method’s usability in the real world, and test it by comparing possible biological conclusions, through a DE analysis experiment. We have used both spline and polynomial regressions strategies as means of assessing differences in biological interpretability. The overall results using each regression technique are displayed in figure 5A and B via volcano plots, the full list of results is available in the Supplementary Material 4). The analysis of these materials elucidates that both regression methods are equivalent which suggests that the spline regression detected a linear pattern in these MA data. This conclusion goes in line with descriptions in the literature. According to the characterisation provided by David, E. (Edwards, 2003), the intensity value detected by the MA scanner is equivalent to the actual value of expression multiplied by two constants (equation 7). Through the proposed mathematical equation, the error at probe and sample levels are considered in a linear relationship. Significant differences in the DE results provided by the two methods differed on only two genes: KRTAP20-2 (Keratin-associated protein 20-2) and MT-ND1 (NADH-ubiquinone oxidoreductase chain 1); these differences are significant in polynomial regression, but not in the case of spline.

**Fig. 5.**
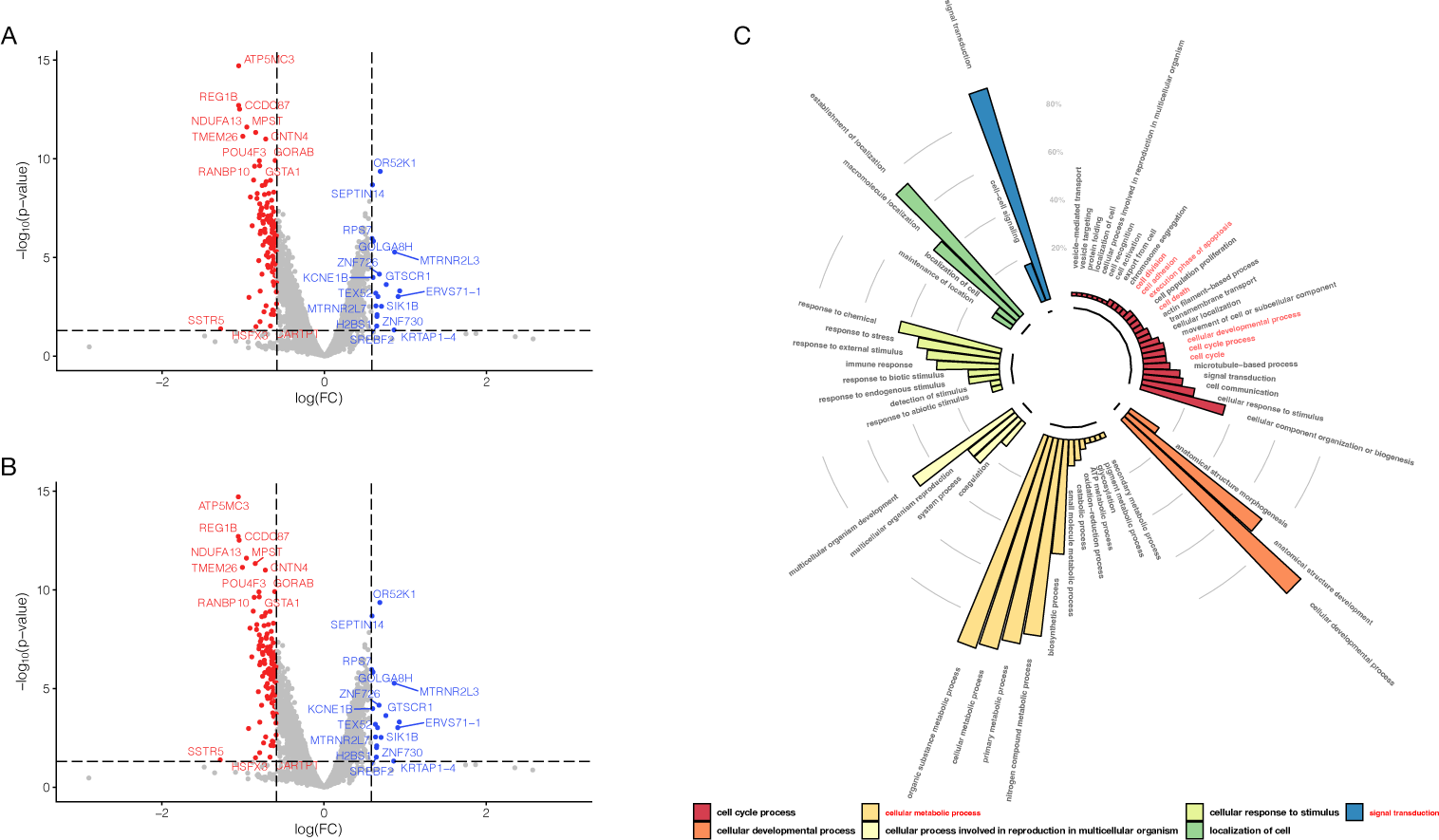
DE analysis results using spline and polynomial regressions. A, B. Volcano plots of the DE using both regression techniques. C. Enrichment analysis results using the polynomial dataset (where biological processes found in other works in literature are highlighted in red). In A and B, the dots in red are under-expressed genes and the dots in blue are over-expressed genes (prostate cancer vs. control). In C, colours represent the first rank of GO biological processes, while each column refers to the second rank; the axis reflects the percentage of DE genes in the list associated with a particular annotation term.

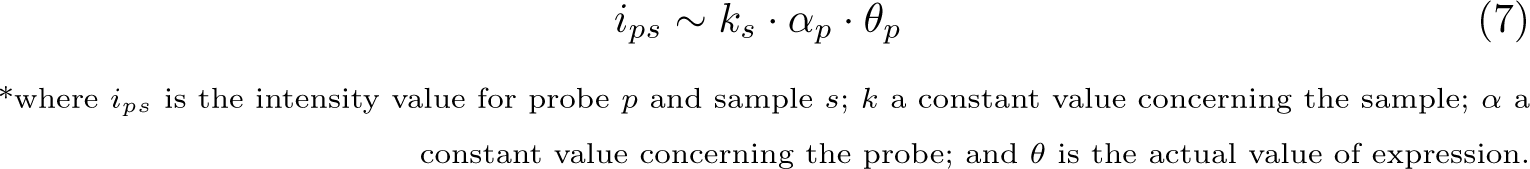

To compare our results with those found in the literature, we ran a gene enrichment experiment against Gene Ontology (GO) (Carbon et al., 2021) biological processes using PANTHER (Mi et al., 2021). Given the theoretical linear behaviour of MA data discussed above, we decided to take polynomial regression as the technique for further steps. Results are presented in figure 5C. Here, biological processes found in other publications on Prostate Cancer are highlighted in red. (Yang et al., 2014) reported differences in biological adhesion, while (Fan et al., 2018) described changes in cell cycle control, cell death regulation, cell development, and focal adhesion. (Sun et al., 2019) and (Verma et al., 2019) reported alterations in the hippo signalling pathway, negative regulation of growth, and multiple metabolic progression.

### Generation of the Real-World Data repository

#### Mining of Demographic Features

The demographical characterisation of patient populations is of utmost importance in the context of clinical trials. The overall outcome of a clinical intervention may differ significantly depending on age (Montes et al., 2018; Przemska-Kosicka et al., 2018), biological sex (Basoglu and Tasbakan, 2018; Gabler et al., 2012) or ethnical background (Morris et al., 2019; Kopel et al., 2020). Additionally, in the case of drug studies, information about body weight is also crucial for the analysis/simulation of pharmacokinetics (Hebbes and Thompson, 2018; Morrish et al., 2011; Smit et al., 2018) and to better understand other unexpected effects (e.g. “BMI paradox” (Tittl et al., 2018; Donini et al., 2020) – in which, patients with extreme weight seem to exhibit better clinical outcomes in metabolic-related diseases). Consequently, we mined GEO for information on five clinical variables, namely age, Body Mass Index (BMI), height, and ethnicity. Given the unstructured nature of GEO metadata, few samples are fully described according to these features, which required the creation of different datasets. Furthermore, demographic features were often embedded in natural text – which led us to apply Natural Language Processing (NLP) techniques to extract and organise them. Figure 6 presents a graphical description of the obtained GEO patient populations. Visual analysis of the data suggests that populations seem to be skewed towards a Western representation of demographics. This fact was highlighted by the ethnological dispersion (figure 6D) of patients, supported by values consisting of age, BMI, and height (figure 6A, B, and C, respectively). However, the comparison against the reference demographics of these populations was not significant (results from these statistical tests can be found in Supplementary Table 5). As such, despite being able to simulate a pool of patients (following predetermined demographic distributions), we cannot extract prevalence data given that the global population of samples does not follow what we observe in reference to regional populations.

**Fig. 6.**
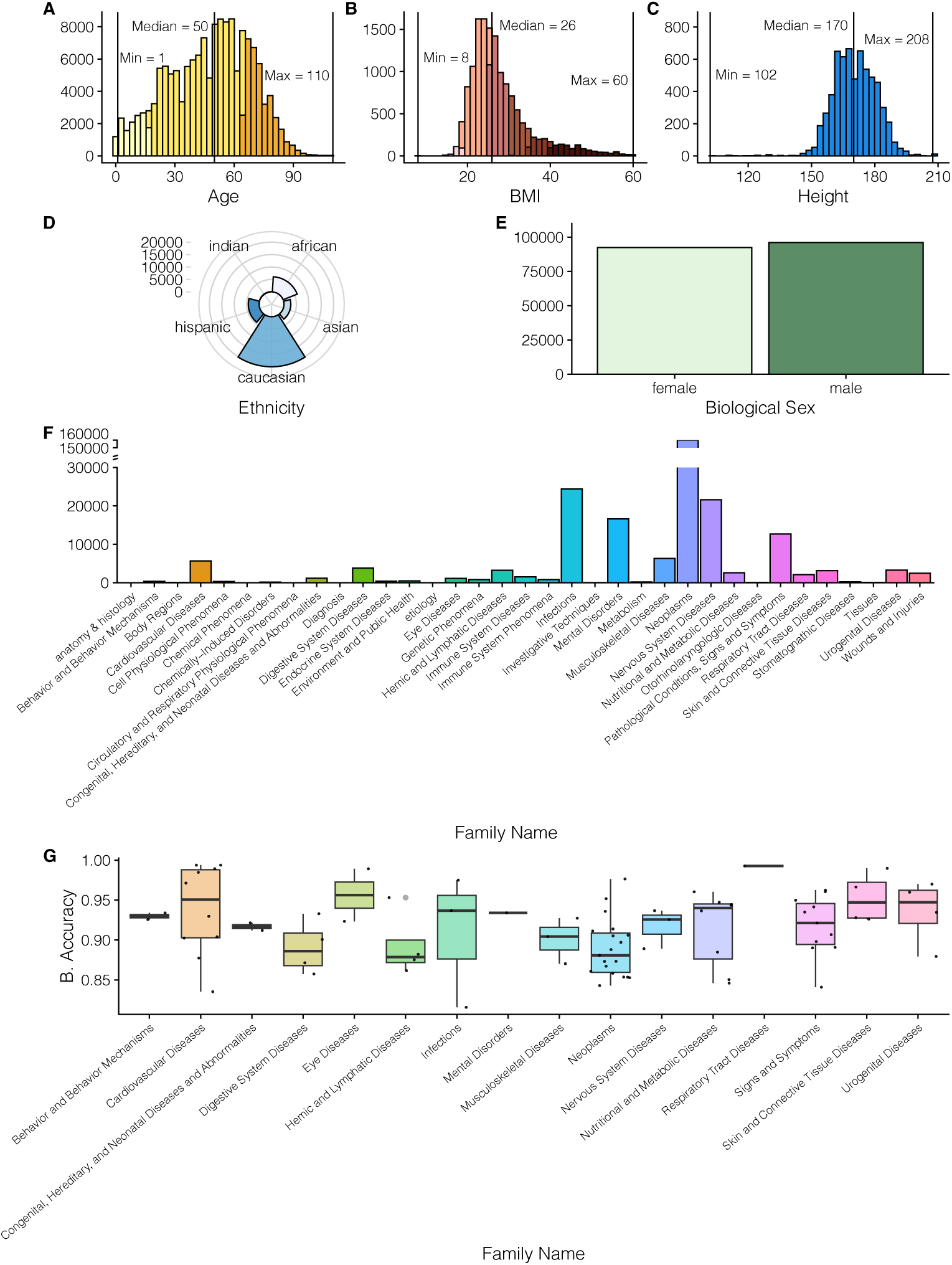
Demographic description of MA samples in GEO. A, Age distribution of samples (n=152,237). Colours represent various age groupings: light yellow – childhood; yellow – adulthood; orange – elderhood. B, BMI distribution of samples (n=14,243). Different shades of red represent different gradings of GEO: very light – underweight; light – normal; red – pre-obese; dark– obese class I; very dark – obese class II. C, Height distribution of samples (n=6,200). D, Ethnicity background ratios (n=33,465). E, Biological sex ratio of samples (n=188,618). F, MeSH terms of disease conditions (top leaf in the MeSH ontology tree) found in GEO samples. G, balanced accuracy of the ML_clinical_ models aimed at predicting samples’ medical conditions. Results are grouped by the MeSH family. Each black dot corresponds to the average balanced accuracy of a medical condition in that family.

Mining demographical metadata from GEO samples is not a new subject. Other works have accomplished this task either by manual curation (STARGEO (Hadley et al., 2017), and OMiCC (Shah et al., 2016)) or by applying AI and NLP techniques (MetaSRA (Bernstein et al., 2017) and ALE (Giles et al., 2017)). However, these works disregard some parts of the demographical description by exclusively reporting the age of the patient and, exclusively in the case of ALE, gender. Additionally, the methods applying manual curation are restricted to the samples presently included in their database, which excludes its usage in later works. Further differences among these tools were discussed by (Wang et al., 2019) in their review of the subject. In our dataset, we have gone a bit further than current methods in the literature by exploring a larger number of demographical features. Furthermore, our pipeline considers the various forms under which demographical information is conveyed in GEO. Our text-processing methodology standardizes synonyms and units (e.g., age values, that may come in weeks, months or years were converted to years in cases; an array of available gender notations were converted to “male” and “female”) which allows full machine interpretability and integration.

#### Mining of Medical Conditions

A similar strategy was used to mine for information on medical conditions associated with the patient samples. The title of each GEO-GSE experiment study and the accompanying text (present in GEO samples’ metadata) was used to assign mentioned medical conditions and their status as healthy, control, or intervention case. To use this RWD repository in a TPMS framework that relies on the Biological Effectors Database (BED) (Jorba et al., 2020), which maps medical conditions to their protein effectors, we mined metadata for the medical terms in BED (300 conditions at the time of analysis). BED links biological processes (adverse drug events, drug indications, medical conditions) with their associated proteins, based on literature reviews that list genes and proteins related to specific conditions and their roles in disease pathways. Extracted pathological attributes in GEO samples were mapped to the BED condition terms. This process was evaluated by manually checking if BED terms were correctly assigned, attaining a median accuracy of 95.33% (detailed accuracy per medical condition is displayed in Supplementary Table 6). However, from the 300 total terms only a portion had corresponding samples in GEO, leading to the final total 162 tagged medical conditions. Regardless, linking these terms to BED permitted associating each medical condition and consequently each sample in GEO to literature-derived molecular descriptions of disease. Figure 6F describes the overall distribution of GEO patients across Medical Subject Headings (MeSH) (for Biotechnology Information and of Medicine, 2021) conditions groups. As in the case of the demographical description, the distribution of diseases highlights the fact that patient samples in GEO reflect the interests of the scientific community and corresponding funding agencies. Most common medical conditions correspond to those mostly investigated (e.g., cancer, neurological and psychological diseases), and not to those more common in societies.

#### Prediction of Medical Conditions

Commonly, clinical annotation in GEO is restricted to medical conditions concerning the experiments’ scope. To further complement medical condition annotation on the dataset, we resorted to ML classifiers – that we will refer as ML_clinical_ moving forward. The input architecture of these models was defined by the BED medical condition to be predicted. Features were defined by BED information on disease-related proteins/-genes. Specifically, genes/proteins annotated as being linked to the models’ phenotype defined what information is taken as input by each ML_clinical_ model. However, due to the different MA platform coverage displayed by the samples – a subject discussed in previous sections of this work, various models were trained corresponding to the same BED medical condition to be able to counteract the large number of missing probes. A global depiction of the CV Ballanced Accuracy (BACC) (Equation 8) metrics is displayed in figure 6G. Here, given the large number of BED conditions, different pathologies are grouped into MeSH condition groups – as in Figure 6F; for further detail, extensive results are available in Supplementary Table 7. All ML_clinical_ models attaining over 80% CV-BACC were kept, and sequentially used (sorted in descending order according to their BACC) to classify MA samples in the RWD repository. Across all available BED diseases, ML_clinical_ models reached at least 81.57% balanced accuracy (average: 91.47%; standard deviation: 0.044). However, we were unable to produce viable ML_clinical_ models for 59.02% of BED medical conditions mainly due to a lack of GEO samples from those specific pathologies. Regardless, the overall accuracy obtained through these models proves MACAROON’s suitability for ML experiments (in addition to the DE analysis presented and discussed in previous sections). Furthermore, these models will also be used for filling the gaps in missing metadata, as well as studying possible comorbidities.

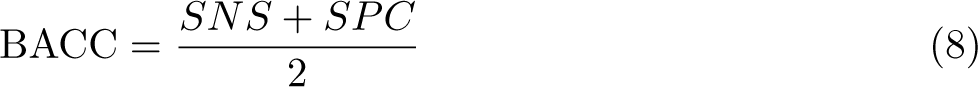

## 3 Conclusion

In this work, we presented a global pipeline for the usage of the GEO database as a source of RWD data. Our global goal was to link demographical and clinical information (derived from GEO samples metadata) with the patients’ underlying molecular patterns (derived from MA data profiles). Our approach begins with an acknowledgment of the predominance of processed data within GEO. Our methodology revolves around the integration of MA data from diverse sources, encompassing various MA platforms, into a unified dataset. Our emphasis lies in the ability to incorporate samples from past experiments, particularly in scenarios where raw intensity values are absent, and investigators only supply processed and normalised files. Through this concerted effort, we aim to streamline data management and improve the comprehensiveness of MA datasets within the GEO repository. The different processing techniques for the normalisation of MA data, and the fact that often supplementary metadata lacks precise information, negatively influence the standardisation process and the conduction of data integration for Big Data experiments. To overcome these challenges, our processing protocol consists of two main steps: (1) samples metadata enrichment using a natural-language processing algorithm; and (2) HK-based normalisation using cross-GEO background datasets.

We have tested each intermediary step independently, and the results show great improvement as we have gone through each stage. The usage of MACAROON was tested in a prostate cancer case study and could reproduce the biological conclusions found in previous works from the literature analysing the transcriptome of the study samples. These results support the usage of MACAROON in future transcriptomics experiments as a novel approach for MA data integration and the exploration biological patterns in pre-processed data. Moreover, it runs independently of previous data processing, allowing the usage of bigger and more diverse cohorts – as required for the discovery of new, patient-centred biomarkers and pharmacological targets. MACAROON has been demonstrated to be a good method for integrating data from different MA datasets when compared to other available methods – better depicting the biological biological mechanisms in the original pre-processed datasets.

Additionally, we explored the usage of GEO metadata as a source of RWD. In addition to the medical conditions linked to each sample, we took the guidance of typical information reported in RCTs and mined for demographical information. This information, besides describing the patient population, allows to assess possible clinical nuances dependent on age, biological sex, ethnicity, and body mass distribution. Despite not being able to significantly modulate any predefined human population, which for instance excludes the possibility of prevalence studies, our current data allow us to randomly choose samples belonging to a custom demographical distribution, according to the research purpose and interest. Expectedly, a larger portion of patients’ metadata does not comprise these data. Future steps to this work would indeed include the usage of ML to extract this information from patterns within the MA data itself. Using these methodologies, it will be possible to include further samples in clinical *in silico* experiments and explore an even more diverse set of patients.

## 4 Methods

### MACAROON, transcriptomics data obtention and processing

#### Obtention of GEO data and dataset generation

Transcriptomic datasets were constructed using MA data available in the GEO repository (Barrett et al., 2013). We have accessed all the available MA human samples (as of 22/12/2020) using the provided “Repository browser” tool. Through this process, we obtained a total of 27,173 series. Each GEO series was divided into its different samples, providing 2,017,019 MA expression profiles corresponding to the column “Value” on the sample page. We kept all the data files in their processed form (as made available by the submitting researcher). Moreover, the metadata matrix (included in the “Data table header descriptions” table of the sample page) was also kept and used in subsequent steps. This procedure was performed with Python 3.9.0 using the package Beautiful Soup 4 (Richardson, 2007). We obtained a total of 1,866,292 samples containing processed MA data files, 211,384 of these did not contain any expression data (were empty entries in the database).

As probe names are platform-dependent (different platforms use different ids for the same genes), probe ids were mapped to their corresponding gene id. The soft file (”GPLXXX family.soft.gz”) corresponding to the platform mentioned in the samples’ metadata was downloaded and processed. To account for “custom platforms” and maximize speed, a local cross-GEO database including platform-specific ids and the corresponding Entrez ID was constructed, and expression files were reindexed according to it. Genes that included multiple intensity measurements were summarized and their mean intensity was kept. To maximize compatibility with the BED, probes were also mapped to the corresponding Uniprot ID.

#### MA integration (postprocessing and normalisation)

##### Composition of datasets

A series of expression-related features were extracted from the MA datasets mentioned in the previous steps. Statistical descriptions such as: the maximum value, minimum value, average, standard deviation, and variation was extracted. Additionally, the value corresponding to the expression of five HK genes was kept. HK genes are responsible for the basic functions of the cell (Butte et al., 2001) and are universally expressed in all tissues and cell types (Zhu et al., 2008). Given their known variation in the expression levels in some or conditions (Greer et al., 2010), we ranked known HK genes according to the standard deviation of their expression across multiple tissues, as published by (Eisenberg and Levanon, 2013). Additionally, as means of maximizing consensus, the ranked list was filtered to only include genes appearing in a minimum of 13 different previously available lists, as reported by Zhang et al. (2015) (Zhang et al., 2015). The final HK genes list is available in Table 3.

##### Inference of pre-processing transformations

###### Text mining

The metadata description of the MA samples was processed using a text mining pipeline to build datasets that would allow the training of ML_processing_ models to predict pre-processing procedures applied to the samples. Given the complexity of some of the sentences, descriptions comprising more than two phrases were excluded. With the remaining examples, we queried the samples for “hot words”: “calibrat”, which is a simplified word that allows the detection of various verbal forms of calibration; “intensity” ; “log” ; “log 10” and “log10” ; “log 2” and “log2”, “mas”, “normali”, a simplified word that allows the detection of American and British English, as well as various verbal forms normalisation, “quantile”, “ratio”, “raw” ; and “rma”. The resulting dataset included samples with one or more “hot words”, as portrayed in table 4. A random selection of 5000 samples, for each of the 6 groups defined in table 4 was recorded and joined with the expression-related features previously described to compose the datasets used for predicting pre-processing of samples (sections below), 20% of the samples were reserved for benchmarking of the trained models.

###### ML models for pre-processing inference

*Model Architecture.* The datasets described in the methods’ sections above were used to train ML_processing_ classifiers to predict what methods had been applied as data pre-processing when these details were unknown. We used a set of different ML algorithms: Decision tree (Breiman et al., 2017), Elastic net (Zou and Hastie, 2005), Elastic net with threshold optimization (Boughorbel et al., 2017), Generalized linear model (Madsen and Thyregod, 2010), Linear classifier (Fukunaga, 1990), Multi-layer perceptron (Haykin, 1998), Naive Bayes Classifier (Russell and Norvig, 2010), Quadratic classifier (Haykin, 1998), and Support Vector Machines (Ben-Hur et al., 2008). Additionally, in each of the models, we also carried out a feature selection process based upon a diversified set of methods described in Supplementary Table 8. Most models were included in MATLAB (version 9.2) (Inc., 2017) and its “Statistics and Machine Learning Toolbox” (Henson and Cetto, 2005). Models that were not available (Linear and Quadratic classifiers, and threshold optimization) were deployed using custom MATLAB code. The hyperparameters of the applied models are described in table 5. The threshold optimization step in Elastic net with threshold optimization models was done by calculation of the Receiver-Operator Curve (ROC). The final models used for the prediction of MA samples’ pre-processing consists of soft-voting ensemble models comprising the top-three best-performing models in each case/dataset.

**Table 5.** ML models hyperparameters. Parameters not mentioned in this table were left as default.

| Model | Hyperparameters | Model | Hyperparameters |
| --- | --- | --- | --- |
| *Decision tree* | Prune ratio: 0; Minimum samples per leaf: 1; | *Multi-layer perceptron* | No hidden layers: 1; No hidden nodes: 2; No epochs: 200; Activation function: Tangent sigmoid; |
| *Elastic net* | Alpha = 0.01 | *Naïve Bayes Classifier* | - |
| *Generalized linear model* | Distribution: binomial; Link: Normal inverse; | *Support Vector Machines* | - |

Validation of pre-processing inference ML_processing_ models was performed by splitting the dataset (defined in sections above and according to table 4) into training and test sets. 20% of samples were reserved for benchmarking, and the remaining 80% were used for training. Training was done in a 25-k cross-validation strategy. The models showing the best training metrics were kept and used in the remaining steps.

*Prediction Confidence.* To improve the ML_processing_ models’ confidence, the raw score (model output before the output activation functions) from each model in the various ensembles was recorded. Scores were split into 5%-quantile intervals and the mean accuracy was calculated for each of them. This value was used to check the models’ certainty level depending on the raw score value. Low-confidence regions were disregarded and only predictions falling into intervals with more than 95% mean accuracy were considered (supplementary figure 1).

###### Uniformization of processing techniques

Samples were classified by ML_processing_ models. In cases where the original preprocessing annotation was unavailable or unclear, expression values were transformed according to classification results. The transformations applied are described in equations 9 and 10.

Where log2 is detected,

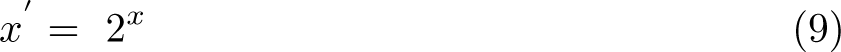

And, where log10 is detected,

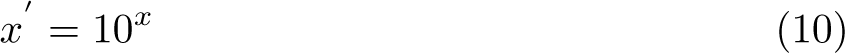

###### Cross-sample normalisation method

After undoing logarithm transformations, we applied a normalisation technique to uniformize all samples in the repository. Given that different samples may have been acquired in different settings (e.g., different MA scanners, processing software), and that we could not resort to control measurements in the samples (e.g., control probes), we applied a normalisation strategy that would level the various samples using a biologically intrinsic value in the samples. We have used the intensity values of HK genes to infer a sample-specific normalisation curve that would be capable of normalizing the expression of the remaining genes. The intensity values from the least variable 40 HK genes (in Table 3) were statistically described across all samples in the repository. Given that none of the genes shows a normal distribution of intensity, we have used a cross-GEO background dataset containing the median expression values of these 40 genes (described in Supplementary Table 9). For each sample, we built a dataset in which the median HK expression values were mapped to the expression value in the given sample. This sample-specific dataset was used to train two types of regressors – (1) spline (Inc., 2017) and (2) polynomial regressors (Inc., 2017) – that were applied to the remaining genes in the sample dataset. As results from both regressions were equivalent, future steps were performed using the polynomial regression for its stronger theoretical support.

###### Benchmark of MACAROON against other integration methods

We have tested MACAROON against a YuGene (Cao et al., 2014) as means of evaluating its performance in comparison to an original MA dataset (GEO series GSE102124 (Sowalsky et al., 2018) – corresponding to the “Gene expression profiling of treated and untreated primary prostate cancer”). The three datasets were submitted to a DE analysis where expression values between treated and untreated samples were compared. The analysis was conducted in R using the package *limma* v. 3.19 (Ritchie et al., 2015). Overlap analysis was performed with the top-50 and bottom-50 genes from each dataset in respect to log FC.

###### Comparative case study

To evaluate the performance of MACAROON, we collected 200 samples of Prostate Cancer from 63 GEO series (see Supplementary Table 3 for further detail). To account for possible errors in the ML-based stages of MACAROON, maximum expression values in each sample were assessed, and samples corresponding to outlier maximum values were excluded (applying the interquartile range criterion (Dekking et al., 2005)). DE analysis resulted from the comparison between intervention vs. control samples in the corresponding GEO series. DEGs were selected using a threshold *|*logFC*|>*=0.5 and FDR *<*0.05, using a Student’s T-test with Bonferroni correction. DEGs were analysed for Gene Ontology BP (Carbon et al., 2021) enrichment using PANTHER (Mi et al., 2021).

### Metadata mining and RWD acquisition

In order to accomplish the second objective of this work – to build an RWD repository from GEO datasets – we mined metadata information provided alongside GEO samples. Medical conditions and demographical information were extracted from each entry. Additionally, knowing the significant differences existing between gene expression across tissues, samples’ tissue information was also retrieved. Information was mined from the text descriptions of GEO samples.

#### Demographical features

To assign the samples’ demographic characteristics, we again mined the provided metadata text for information on age, biological sex, ethnicity, BMI and height. Age was extracted mainly from the subfields “age” and “birth”. The obtained values were then standardized to comprise the age of the subject in years. Biological sex was mined from the subfield “gender”. Different available classes (given that sex is described using different notations) were standardized to “female” and “male”. Ethnicity was obtained from the fields “ethnicity”, “race” and “population”. The identified vocabulary was standardized to “african”, “asiatic”, “caucasian”, “hispanic” and “indian”. Information on BMI was mined from the subfield “BMI”, with no further transformations. Height values were extracted from the subfield “height”. Units were mined from text and converted to centimetres (when values were presented in other units).

#### Medical conditions

Samples were tagged according to the medical condition provided in GEO series and the samples’ titles and descriptions. Medical conditions, corresponding to BED terms, were assigned to each sample by using a manually curated synonym dictionary where metadata sentences were mined for a series of hot words. Additionally, healthy and control samples were explicitly mined to account for different experimental designs in the GEO series. The different BED terms were mapped to MeSH terms (for Biotechnology Information and of Medicine, 2021) using MetaMap (Bodenreider, 2004) – a tool to map biomedical terms referred to in raw text to controlled terms in the Unified Medical Language System (UMLS) and corresponding Condition ID (CUI). This allowed the hierarchical grouping of the different medical conditions for grouped analysis.

#### Information on source tissue

To account for expression differences, information regarding the samples’ body tissues was also extracted from the metadata file. To reduce redundancy, and to allow for hierarchical classification of body tissues, terms were mapped to the Brenda Tissue Ontology (Gremse et al., 2011).

#### ML models to predict BED medical conditions

For each BED medical condition mined from the GEO samples, we trained a set of ML_clinical_ models aimed at deciphering further medical conditions in the samples. Proteins associated with each medical condition in the BED were selected as features.

Models were trained in a grid-search fashion, including feature selection (following the strategy applied to ML_processing_). A maximum of 2,000 samples were used for each class, with the positive class containing GEO samples from which the specific BED medical condition was mined. On the other side, the negative class corresponded to a random set of samples where that medical condition was not mentioned. To account for variability in the number of samples, models were selected based on their cross-validation (k=10) BACC, where only the models with balanced accuracy above 80% (p-value*<*0.01) were kept. To account for missing values, for each medical condition, models had a different number of features (ranging from 2 to 5).

**Supplementary information.** This article has accompanying supplementary files.

## Declarations

- Funding: PMF, JMGI, GJ and JF are employees of Anaxomics Biotech S.L. JMM is an employee and co-founder of Anaxomics Biotech S.L. BO is a scientific advisor for Anaxomics Biotech but does not perceive any funding support for these services. PMF and JMGI received funding from the European Union’s Horizon 2020 research and innovation programme under the Marie Sklodowska-Curie grant agreement No 860303 (proEVLifeCycle) and 859962 (SECRET), respectively.
- Conflict of interest/Competing interests: None declared.
- Ethics approval and consent to participate: The patient data included in this work have been obtained from GEO database, where cell lines and other sources of information not adequate for our proposes were excluded. GEO is a public repository that contains molecular data that could be considered sensible from certain ethical perspectives given that these data describe the molecular status of patients. However, this source of data does not contain personal details or any other way to allow the patient identification and neither other clinical detail of them. Despite this fact and to reinforce the patients’ privacy, all patients included in this study were submitted to a pseudonymisation protocol to avoid any possible link between our patient repository and the original GEO identification. This data treatment has been done according to the EU regulation on personal data and its management (specifically, GDPR – EU regulation 2016/679). Additionally, we also choose not to provide our patient repository to avoid any possible misuse by third people or entities about the clinical information estimated during our analyses and to ensure the application of the “appropriate safeguards” according to this regulation. All measures taken in this study are driven to the aim pursued, respect the essence of the right to data protection and provide for suitable and specific measures to safeguard the fundamental rights and the interests of the data subject. The consent of the participants has been treated according to the exposed regulations and data management and considering that this study is a non-interventional work with scientific and public interest, based on historical research and with statistical purposes that show aggregated information to extract the biological and clinical conclusions.

- Consent for publication: Not applicable.
- Data availability: The datasets supporting the conclusions of this article are publicly available in the GEO repository, the whole database is available at https://ftp.ncbi. nlm.nih.gov/geo/. Series applied in benchmarking and case studies are referenced in the methods and references sections, and in the supplementary material.
- Materials availability: Not applicable.
- Code availability: Not applicable.
- Author contribution: PMF, methodology, software, validation, formal analysis, investigation, resources, Writing – original draft, Writing – review & editing, visualisation. JMGI, methodology, data curation, formal analysis, Writing – review & editing. GJ, conceptualization, methodology, Validation, formal analysis, data curation, Writing – review & editing. BO, Supervision, Writing – review & editing. JF, Conceptualization, Funding acquisition, Investigation, Project administration, Supervision, Validation, Writing – review & editing. JMM, Conceptualization, Formal analysis, Funding acquisition, Investigation, Methodology, Project administration, Resources, Software, Supervision, Validation, Visualization, Writing – original draft, Writing – review & editing.

## Footnotes

1 The concept of dynamic range refers to the ratio between the maximum and minimum value a specific technology or device may withstand.

