## Supplementary Material for "The usage of transcriptomics datasets as sources of Real-World Data for clinical trialling"

### SUPPLEMENTARY FIGURES

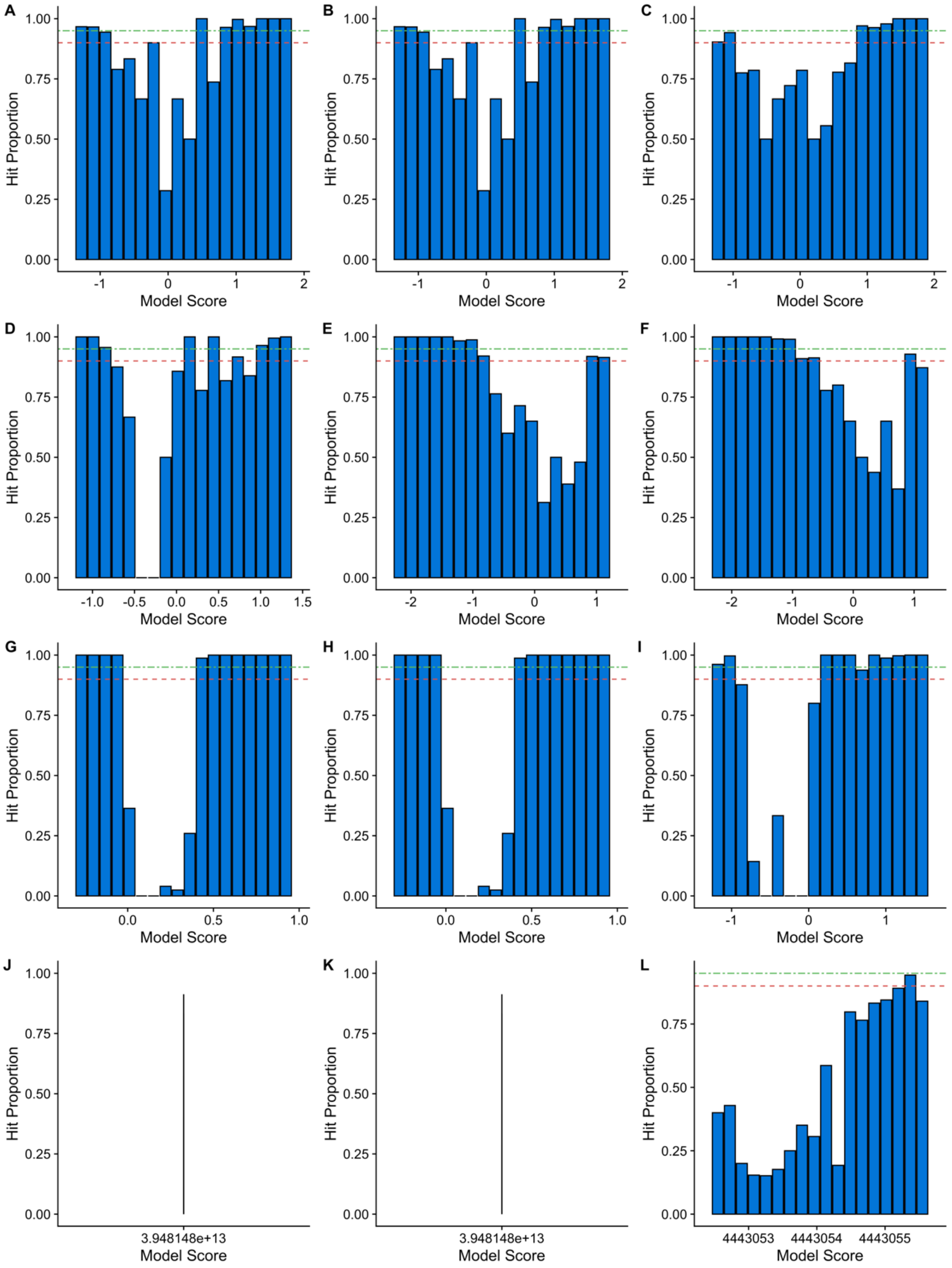

**Supplementary figure 1.** Machine Learning models' output before activation function. In each row, each plot concerns one of the models in the respective ensemble.  A-C: Log2; D-F: Log10; G-I: Log; J-L: Ratio; M-O: Normalization.

### SUPPLEMENTARY TABLES

**Supplementary Table 1.** Test performance of the final sample processing detection ensemble model.

| ***Metric*** | **Log 2** | **Log 10** | **Ratio** |
|---|---|---|---|
| *True Positives (TP)* | 861 | 706 | 895 |
| *True Negatives (TN)* | 649 | 828 | 55 |
| *False Positives (FP)* | 29 | 19 | 5 |
| *False Negatives (FN)* | 23 | 18 | 0 |
| *Accuracy (ACC)* | 96.67 % | 97.64 % | 99.47 % |
| *Precission (PRE)* | 96.74 % | 97.37 % | 99.44 % |
| *Sensitivity (SNS)* | 97.39 % | 97.51 % | 100.00 % |
| *Specificity (SPC)* | 95.72 % | 97.75 % | 91.67 % |
| *MCC* | 93.22 % | 95.26 % | 95.48 % |
| *Balanced ACC* | 96.56 % | 97.64 % | 95.83 % |
| *F1-Score* | 97.07 % | 97.45 % | 99.72 % |
| *AUC* | 96.56 % | 97.64 % | 95.83 % |

**Supplementary Table 2.** Training metrics of the sample processing models.

| ***Metric*** | **Log 2** | **Log 10** | **Ratio** |
|---|---|---|---|
| *True Positives (TP)* | 3635 | 2723 | 3702 |
| *True Negatives (TN)* | 2362 | 3532 | 321 |
| *False Positives (FP)* | 48 | 62 | 22 |
| *False Negatives (FN)* | 51 | 14 | 1 |
| *Accuracy (ACC)* | 98.37 % | 98.79 % | 99.43 % |
| *Precission (PRE)* | 98.69 % | 97.77 % | 99.40 % |
| *Sensitivity (SNS)* | 98.61 % | 99.48 % | 99.97 % |
| *Specificity (SPC)* | 98.00 % | 98.27 % | 93.58 % |
| *MCC* | 96.60 % | 97.57 % | 96.29 % |
| *Balanced ACC* | 98.31 % | 98.88 % | 96.77 % |
| *F1-Score* | 98.65 % | 98.62 % | 99.69 % |
| *AUC* | 98.31 % | 98.88 % | 96.77 % |

**Supplementary Table 3.** Prostate Cancer GEO experiments applied in the DE case study.

| *GEO ID* | Title | Date of Submission | Organism | PMID |
|---|---|---|---|---|
| *GSE100301* | Genome-wide analysis of galectin-4 mediated gene signature in PCa | 21/06/2017 | Homo sapiens | |
| *GSE101982* | Expression data from LNCaP clonal isolates over-expressing HIST1H1A or Vector Control. | 27/07/2017 | Homo sapiens | 29983878 |
| *GSE102124* | Gene expression profiling of treated and untreated primary prostate cancer | 01/08/2017 | Homo sapiens | 29921690 |
| *GSE102616* | ZFX acts as a transcriptional activator in multiple types of human tumors by binding downstream of transcription start sites at the majority of CpG island promoters | 14/08/2017 | Homo sapiens | 29429977 |
| *GSE103864* | Identification of global XBP1s target gene expression in human prostate cancer cells | 14/09/2017 | Homo sapiens | 30679434 |
| *GSE104418* | Examining alterations in gene expression of PC3-wt (PSMA -ve) vs PC3-PSMA (PSMA +ve) cells | 28/09/2017 | Homo sapiens | 29141866 |
| *GSE10585* | Egr1 regulates the coordinated expression of numerous EGF receptor target genes as identified by ChIP-on-chip | 20/02/2008 | Homo sapiens | 19032775 |
| *GSE107245* | Proteotranscriptomic profiling of potential E6AP targets in prostate cancer cells | 21/11/2017 | Homo sapiens | 29463595 |
| *GSE108545* | Transcriptomic Reprogramming of Prostate Cancer Cells Driven by Stroma-Derived SPINK1 | 26/12/2017 | Homo sapiens | 30333494 |
| *GSE111648* | Microbial signatures in prostate cancer | 09/03/2018 | Homo sapiens | 32076488 |
| *GSE113234* | Circulating microRNAs combined with PSA for an accurate and non-invasive prostate cancer detection | 17/04/2018 | Homo sapiens | 30452625 |
| *GSE113309* | BMI1 drives metastasis of prostate cancer in Caucasian and African-American men and is a potential therapeutic target: hypothesis | 18/04/2018 | Homo sapiens | |

| | | | | |
|---|---|---|---|---|
| | tested in race-specific models | | | |
| *GSE114737* | Optimized ChIP-seq procedure facilitates hormone receptor profiling in human tumors | 21/05/2018 | Homo sapiens | 30620009 |
| *GSE115615* | BRG1 occupancy profiling by high throughput sequencing from control and PTEN knockdown 22RV-1 cells | 11/06/2018 | Homo sapiens | 30496141 |
| *GSE115619* | SWI/SNF Chromatin Remodeling Factor BRG1 is Synthetic Required for PTEN-deficient Prostate Cancer | 11/06/2018 | Homo sapiens | 30496141 |
| *GSE118959* | J_093015-circRNA-33 AS-CR-005 Human CircRNA V2 microarray 01272016 | 23/08/2018 | Homo sapiens | 31341219 |
| *GSE120738* | Integrative epigenetic taxonomy of primary prostate cancer [ChIP-Seq] | 02/10/2018 | Homo sapiens | 30464211 |
| *GSE120742* | Integrative epigenetic taxonomy of primary prostate cancer | 02/10/2018 | Homo sapiens | 30464211 |
| *GSE129739* | Gene expression-based, whole genome-wide expression profile differences for CUL4B-depleted LNCaP cells compared with control | 12/04/2019 | Homo sapiens | 31111526 |
| *GSE14092* | An Integrated Network of Androgen Receptor and TMPRSS2-ERG Gene Fusion in Prostate Cancer Progression (II) | 22/12/2008 | Homo sapiens | 20478527 |
| *GSE14996* | Multisampled Lethal Metastatic Prostate Cancer Copy Number Analysis | 25/02/2009 | Homo sapiens | 19363497 |
| *GSE15792* | Expression profiles in PCARNEQ ncRNA depletion in prostate cancer cells | 23/04/2009 | Homo sapiens | |
| *GSE26022* | [Gene Expression Training Set] Protein-coding and MicroRNA Biomarkers of Recurrence of Prostate Cancer Following Radical Prostatectomy | 11/12/2010 | Homo sapiens | |
| *GSE28241* | Menthol effect on gene expression | 29/03/2011 | Homo sapiens | 22580005 |

| | | | | |
|---|---|---|---|---|
| | profile in malignant prostate cancer cell lines, PC-3 | | | |
| *GSE31728* | Transcriptome sequencing across a prostate cancer cohort identifies PCAT-1, an unannotated lincRNA implicated in disease progression | 29/08/2011 | Homo sapiens | 21804560 |
| *GSE33277* | Castration-Resistant Prostate Cancer (CRPC) | 27/10/2011 | Homo sapiens | 22363599 |
| *GSE34893* | MiR-106b-25 cluster targets caspase 7 and focal adhesion in human prostate cancer | 05/01/2012 | Homo sapiens | |
| *GSE35126* | Effects of Cardiac Glycosides on RNA Expression in Prostate Cancer LNCaP-abl Cells | 16/01/2012 | Homo sapiens | 22396588 |
| *GSE35211* | HIP1 mediates prostate carcinogenesis and progression through STAT3 signalling | 19/01/2012 | Homo sapiens | |
| *GSE36165* | Drug efficacy reprogramming against aggressive human prostate cancer | 29/02/2012 | Homo sapiens | 23600803 |
| *GSE37119* | Identification of target genes of cancer-related microRNAs in human cancer | 09/04/2012 | Homo sapiens | 22766839 |
| *GSE38073* | DNA methylation profiling of normal prostates and prostate cancer metastases [Affymetrix] | 21/05/2012 | Homo sapiens | 23345608 |
| *GSE39452* | Expression data of EZH2-dependent genes in prostate cancer cell lines | 18/07/2012 | Homo sapiens | 23239736 |
| *GSE39461* | Role of PRC2 complex components in prostate cancer cell lines | 18/07/2012 | Homo sapiens | 23239736 |
| *GSE42977* | Sequential Binary Gene-Ratio Tests Define a Novel Molecular Diagnostic Strategy for Malignant Pleural Mesothelioma | 18/12/2012 | Homo sapiens | 23493352 |
| *GSE43785* | Redefinition of Human Androgen Responsive Elements [ChIP-Seq, RNA-Seq] | 27/01/2013 | Homo sapiens | 25535248 |
| *GSE45033* | Effect of CCAR1 depletion on androgen-dependent | 12/03/2013 | Homo sapiens | |

| | | | | |
|---|---|---|---|---|
| | gene expression in LNCaP cells | | | |
| *GSE45567* | The transcriptional network for cell cycle regulation in prostate cancer | 27/03/2013 | Homo sapiens | 24239550 |
| *GSE46177* | A DNA hypermethylation profile reveals new potential biomarkers for prostate cancer diagnosis and prognosis | 18/04/2013 | Homo sapiens | |
| *GSE51629* | Treatment of prostate cancer cell lines DU-145 and LNCaP with zebularine | 23/10/2013 | Homo sapiens | 26317032 |
| *GSE52201* | Lysine-Specific Demethylase 1 Has Dual Functions as a Major Regulator of Androgen Receptor Transcriptional Activity | 07/11/2013 | Homo sapiens | 25482560 |
| *GSE53115* | The novel NLR-related protein NWD1 is associated with prostate cancer progression and impacts androgen receptor signaling. | 09/12/2013 | Homo sapiens | 24681825 |
| *GSE55598* | DNA methylation sensitivity at detecting cancer in prostate core biopsies. | 05/03/2014 | Homo sapiens | |
| *GSE55599* | DNA methylation status is more sensitive than gene expression at detecting cancer in prostate core biopsies | 05/03/2014 | Homo sapiens | 24937670 |
| *GSE56243* | Differentially expressed genes after miRNA or siRNA transfection in human cancer cell lines | 26/03/2014 | Homo sapiens | 26325107 |
| *GSE56686* | Quantitation of microRNAs in seminal fluid from men with and without prostate cancer | 10/04/2014 | Homo sapiens | 24859988 |
| *GSE59432* | Effects of SPOP on gene expression in prostate cancer | 15/07/2014 | Homo sapiens | 25274033 |
| *GSE60722* | Regulation of casodex-dependent AR activity by NCOR1 | 25/08/2014 | Homo sapiens | 26968201 |
| *GSE61838* | Genome-wide analysis of full-length Androgen Receptor (AR) and AR Splice Variant (ARv567es) cistromes | 29/09/2014 | Homo sapiens | 25908785 |
| *GSE62610* | High miR-449b expression in prostate | 22/10/2014 | Homo sapiens | 25416653 |

| | | | | |
|---|---|---|---|---|
| | cancer is associated with biochemical recurrence after radical prostatectomy | | | |
| *GSE64528* | Assembly of methylated LSD1 and CHD1 drives AR-dependent transcription and translocation [ChIP-Seq] | 29/12/2014 | Homo sapiens | 26751641 |
| *GSE64530* | Assembly of methylated LSD1 and CHD1 drives AR-dependent transcription and translocation | 29/12/2014 | Homo sapiens | 26751641 |
| *GSE66335* | EphA6 promotes angiogenesis and prostate cancer metastasis and is associated with human prostate cancer progression | 26/02/2015 | Homo sapiens | |
| *GSE67457* | Expression data from prostate cancer cell lines LNCap (androgen dependent) and DU145 (androgen independent), transfected with Pin1 or control siRNA | 31/03/2015 | Homo sapiens | 26039047 |
| *GSE762* | CCFAlmasan_CaP1 | 20/10/2003 | Homo sapiens | |
| *GSE782* | CCF_Almasan_CaP GSE56 | 24/10/2003 | Homo sapiens | |
| *GSE81780* | Contribution of STAT3, IRF1 and PGAM5 to androgen response in prostate cancer cells | 23/05/2016 | Homo sapiens | 28826481 |
| *GSE87481* | Clonal isolates of PC3 cells overexpressing CENPU | 29/09/2016 | Homo sapiens | 27916600 |
| *GSE87482* | Clonal isolates of PC3 cells overexpressing RWDD4 | 29/09/2016 | Homo sapiens | 27916600 |
| *GSE87491* | Analysis of germline modifiers of aggressive prostate cancer using cross-species variation and systems genetics | 29/09/2016 | Homo sapiens | |
| *GSE92574* | Mapping of DHT-responsive or -independent AR-binding sites induced by activated Src in prostate cancer cell lines [RNA-seq] | 19/12/2016 | Homo sapiens | 28055971 |
| *GSE93928* | Altered expression of genes in HOXD-AS1 knockdown prostate cancer cells | 23/01/2017 | Homo sapiens | 28487115 |
| *GSE97549* | Global microarray analysis of | 09/04/2017 | Homo sapiens | 30478421 |

| |
|---|
| ONECUT2 transcription factor overexpression in human prostate cancer cells |

**Supplementary Table 4.** Differential Expression Analysis results using the polynomial regression dataset.

| HUGO Code | Log(FC) | P-Value | DE Status |
|---|---|---|---|
| *SSTR5* | -1.2790 | 0.0414 | DOWN |
| *REG1B* | -1.0570 | 0.0000 | DOWN |
| *ATP5MC3* | -1.0560 | 0.0000 | DOWN |
| *CCDC87* | -1.0454 | 0.0000 | DOWN |
| *TMEM26* | -1.0053 | 0.0000 | DOWN |
| *NDUFA13* | -0.9562 | 0.0000 | DOWN |
| *CCDC187* | -0.9297 | 0.0011 | DOWN |
| *PMM1* | -0.9119 | 0.0000 | DOWN |
| *MARCHF10* | -0.8898 | 0.0000 | DOWN |
| *SCARF1* | -0.8721 | 0.0000 | DOWN |
| *RANBP10* | -0.8601 | 0.0000 | DOWN |
| *MPST* | -0.8471 | 0.0000 | DOWN |
| *HSFX3* | -0.8438 | 0.0321 | DOWN |
| *IL9R* | -0.8315 | 0.0000 | DOWN |
| *USP49* | -0.8282 | 0.0000 | DOWN |
| *C5orf67* | -0.8086 | 0.0000 | DOWN |
| *MMP26* | -0.8022 | 0.0000 | DOWN |
| *POU4F3* | -0.8016 | 0.0000 | DOWN |
| *KRT12* | -0.7994 | 0.0000 | DOWN |
| *GSTA1* | -0.7977 | 0.0000 | DOWN |
| *TBL1Y* | -0.7970 | 0.0000 | DOWN |
| *MT-CO1* | -0.7930 | 0.0178 | DOWN |
| *DCT* | -0.7921 | 0.0000 | DOWN |
| *HSPBAP1* | -0.7912 | 0.0005 | DOWN |
| *C12orf71* | -0.7842 | 0.0000 | DOWN |
| *DTX3* | -0.7832 | 0.0000 | DOWN |
| *PPP6R3* | -0.7725 | 0.0000 | DOWN |
| *SLC51B* | -0.7692 | 0.0001 | DOWN |
| *CPB2* | -0.7664 | 0.0000 | DOWN |
| *LNP1* | -0.7587 | 0.0000 | DOWN |
| *SATB1* | -0.7564 | 0.0000 | DOWN |
| *EGLN1* | -0.7547 | 0.0000 | DOWN |
| *KIF19* | -0.7536 | 0.0000 | DOWN |
| *SLK* | -0.7443 | 0.0057 | DOWN |
| *LSM4* | -0.7383 | 0.0000 | DOWN |
| *MITF* | -0.7348 | 0.0000 | DOWN |
| *FNDC11* | -0.7333 | 0.0000 | DOWN |
| *PRCD* | -0.7309 | 0.0000 | DOWN |
| *GATA6* | -0.7308 | 0.0000 | DOWN |
| *CNTN4* | -0.7230 | 0.0000 | DOWN |
| *TEX51* | -0.7186 | 0.0028 | DOWN |
| *SOS1* | -0.7181 | 0.0000 | DOWN |
| *FAM13B* | -0.7149 | 0.0000 | DOWN |
| *AIRE* | -0.7128 | 0.0000 | DOWN |
| *ADAT1* | -0.7122 | 0.0000 | DOWN |
| *ARHGEF10* | -0.7118 | 0.0000 | DOWN |
| *TMEM268* | -0.7018 | 0.0000 | DOWN |
| *BRPF1* | -0.7006 | 0.0000 | DOWN |
| *SLC26A8* | -0.6978 | 0.0000 | DOWN |
| *APOB* | -0.6964 | 0.0000 | DOWN |
| *IL36B* | -0.6925 | 0.0000 | DOWN |
| *ZNF541* | -0.6912 | 0.0000 | DOWN |
| *KRT15* | -0.6896 | 0.0000 | DOWN |
| *REEP1* | -0.6854 | 0.0000 | DOWN |
| *GPR85* | -0.6805 | 0.0000 | DOWN |
| *VWA5B2* | -0.6798 | 0.0000 | DOWN |
| *ACTR10* | -0.6768 | 0.0000 | DOWN |
| *MZT1* | -0.6763 | 0.0000 | DOWN |
| *CARTPT* | -0.6652 | 0.0298 | DOWN |
| *CHTOP* | -0.6649 | 0.0000 | DOWN |
| *OVCA2* | -0.6615 | 0.0000 | DOWN |
| *DSCC1* | -0.6587 | 0.0000 | DOWN |
| *LRIG2* | -0.6579 | 0.0000 | DOWN |
| *DGKB* | -0.6563 | 0.0000 | DOWN |
| *TSSK6* | -0.6562 | 0.0000 | DOWN |
| *FAM229A* | -0.6544 | 0.0076 | DOWN |
| *ZNF282* | -0.6520 | 0.0000 | DOWN |
| *KLRC3* | -0.6497 | 0.0000 | DOWN |
| *NCOA1* | -0.6479 | 0.0000 | DOWN |
| *HIPK3* | -0.6476 | 0.0000 | DOWN |
| *TOX2* | -0.6451 | 0.0000 | DOWN |
| *PIGA* | -0.6440 | 0.0000 | DOWN |
| *DEFB133* | -0.6435 | 0.0000 | DOWN |
| *UBAC1* | -0.6425 | 0.0001 | DOWN |
| *PHACTR1* | -0.6413 | 0.0000 | DOWN |
| *GPX2* | -0.6394 | 0.0000 | DOWN |
| *CMC2* | -0.6392 | 0.0000 | DOWN |
| *PRR23D2* | -0.6369 | 0.0046 | DOWN |
| *RBM28* | -0.6367 | 0.0000 | DOWN |
| *DDX49* | -0.6328 | 0.0000 | DOWN |
| *LSM14A* | -0.6320 | 0.0000 | DOWN |

| | | | | | | | |
|---|---|---|---|---|---|---|---|
| *BCLAF1* | -0.6312 | 0.0000 | DOWN | *LBX1* | -0.5960 | 0.0000 | DOWN |
| *KDM5A* | -0.6310 | 0.0002 | DOWN | *ARMC3* | -0.5929 | 0.0006 | DOWN |
| *UBASH3A* | -0.6300 | 0.0000 | DOWN | *KLHDC4* | -0.5916 | 0.0000 | DOWN |
| *MUC19* | -0.6280 | 0.0049 | DOWN | *BRD7* | -0.5901 | 0.0000 | DOWN |
| *ADNP* | -0.6255 | 0.0000 | DOWN | *FEV* | -0.5892 | 0.0022 | DOWN |
| *ARFGEF2* | -0.6237 | 0.0000 | DOWN | *HAUS4* | -0.5882 | 0.0002 | DOWN |
| *MLLT10* | -0.6229 | 0.0079 | DOWN | *DST* | -0.5880 | 0.0000 | DOWN |
| *NUP205* | -0.6224 | 0.0000 | DOWN | *SLURP1* | -0.5861 | 0.0000 | DOWN |
| *ANGPTL3* | -0.6200 | 0.0002 | DOWN | *RPS7* | 0.5879 | 0.0000 | UP |
| *UBTFL1* | -0.6193 | 0.0003 | DOWN | *SEPTIN14* | 0.5931 | 0.0000 | UP |
| *KIF23* | -0.6170 | 0.0000 | DOWN | *KCNE1B* | 0.6002 | 0.0001 | UP |
| *DNAJB12* | -0.6163 | 0.0000 | DOWN | *GOLGA8H* | 0.6051 | 0.0000 | UP |
| *INKA2* | -0.6149 | 0.0000 | DOWN | *MTRNR2L6* | 0.6337 | 0.0006 | UP |
| *MEX3A* | -0.6119 | 0.0000 | DOWN | *MTRNR2L7* | 0.6358 | 0.0029 | UP |
| *ARFGAP2* | -0.6095 | 0.0000 | DOWN | *SREBF2* | 0.6457 | 0.0298 | UP |
| *OXTR* | -0.6082 | 0.0001 | DOWN | *H2BS1* | 0.6486 | 0.0098 | UP |
| *GORAB* | -0.6073 | 0.0000 | DOWN | *ZNF730* | 0.6498 | 0.0080 | UP |
| *ARIH1* | -0.6064 | 0.0000 | DOWN | *TEX52* | 0.6606 | 0.0010 | UP |
| *CLIC3* | -0.6058 | 0.0000 | DOWN | *ZNF726* | 0.6789 | 0.0001 | UP |
| *KCNMB4* | -0.6040 | 0.0000 | DOWN | *OR52K1* | 0.6891 | 0.0000 | UP |
| *SPP2* | -0.6036 | 0.0000 | DOWN | *SIK1B* | 0.7045 | 0.0030 | UP |
| *DLAT* | -0.6026 | 0.0000 | DOWN | *GTSCR1* | 0.7631 | 0.0002 | UP |
| *CCR10* | -0.5994 | 0.0000 | DOWN | *KRTAP1-4* | 0.8596 | 0.0473 | UP |
| *DAAM2* | -0.5972 | 0.0000 | DOWN | *MTRNR2L3* | 0.8660 | 0.0000 | UP |
| *ZNF619* | -0.5967 | 0.0000 | DOWN | *ERVS71-1* | 0.9099 | 0.0010 | UP |
| *VAMP4* | -0.5964 | 0.0000 | DOWN | *MEIKIN* | 0.9291 | 0.0005 | UP |

**Supplementary Table 5.** Comparison of obtained demographics information vs. reference populations. Statistical test changes according to the demographics feature. Age: non-parametric comparison of the median (Mann-Whitney), ref. population – median value; Biological-sex: $\chi^2$-squared test of proportions, ref. population – the proportion of female individuals; BMI and height: parametric comparison of means ($\tau$ of Student); ref. population – mean value.

| *Demographical Feature* | Continent | Year | Ref. Population | p-value | Accept $H_0$ |
|---|---|---|---|---|---|
| *Age* | Australia and New Zealand | 2020 | 37.90 | 0.00 | No |
| *Age* | Central Asia | 2020 | 27.06 | 0.00 | No |
| *Age* | Eastern Asia | 2020 | 40.05 | 0.00 | No |
| *Age* | Eastern Europe | 2020 | 41.63 | 0.00 | No |
| *Age* | Global | 2020 | 29.91 | 0.00 | No |
| *Age* | Latin America and the Caribbean | 2020 | 32.25 | 0.00 | No |
| *Age* | Melanesia | 2020 | 25.70 | 0.00 | No |
| *Age* | Micronesia | 2020 | 25.87 | 0.00 | No |
| *Age* | Northern Africa | 2020 | 27.69 | 0.00 | No |
| *Age* | Northern America | 2020 | 39.85 | 0.00 | No |
| *Age* | Northern Europe | 2020 | 41.27 | 0.00 | No |
| *Age* | Polynesia | 2020 | 25.67 | 0.00 | No |
| *Age* | South-eastern Asia | 2020 | 29.91 | 0.00 | No |
| *Age* | Southern Asia | 2020 | 27.61 | 0.00 | No |
| *Age* | Southern Europe | 2020 | 42.98 | 0.00 | No |
| *Age* | Sub-Saharan Africa | 2020 | 20.58 | 0.00 | No |
| *Age* | Western Asia | 2020 | 29.87 | 0.00 | No |
| *Age* | Western Europe | 2020 | 42.97 | 0.00 | No |
| *Biological sex* | Australia and New Zealand | 2020 | 50.53 | 0.00 | No |
| *Biological sex* | Central Asia | 2020 | 50.50 | 0.00 | No |
| *Biological sex* | Eastern Asia | 2020 | 51.10 | 0.00 | No |
| *Biological sex* | Eastern Europe | 2020 | 52.17 | 0.00 | No |
| *Biological sex* | Global | 2020 | 49.95 | 0.00 | No |
| *Biological sex* | Latin America and the Caribbean | 2020 | 50.83 | 0.00 | No |
| *Biological sex* | Melanesia | 2020 | 49.30 | 0.02 | No |
| *Biological sex* | Micronesia | 2020 | 49.84 | 0.00 | No |
| *Biological sex* | Northern Africa | 2020 | 49.88 | 0.00 | No |
| *Biological sex* | Northern America | 2020 | 50.44 | 0.00 | No |
| *Biological sex* | Northern Europe | 2020 | 51.13 | 0.00 | No |
| *Biological sex* | Polynesia | 2020 | 49.18 | 0.48 | Yes |
| *Biological sex* | South-eastern Asia | 2020 | 49.77 | 0.00 | No |
| *Biological sex* | Southern Asia | 2020 | 48.21 | 0.00 | No |
| *Biological sex* | Southern Europe | 2020 | 50.77 | 0.00 | No |
| *Biological sex* | Sub-Saharan Africa | 2020 | 50.12 | 0.00 | No |
| *Biological sex* | Western Asia | 2020 | 44.97 | 0.00 | No |
| *Biological sex* | Western Europe | 2020 | 50.47 | 0.00 | No |
| *BMI* | Australia and New Zealand | 2016 | 27.75 | 0.69 | Yes |
| *BMI* | Central Asia | 2016 | 26.69 | 0.00 | No |
| *BMI* | Eastern Asia | 2016 | 24.27 | 0.00 | No |

| | | | | | |
|---|---|---|---|---|---|
| *BMI* | Eastern Europe | 2016 | 26.92 | 0.00 | No |
| *BMI* | Global | 2016 | 26.75 | 0.00 | No |
| *BMI* | Latin America and the Caribbean | 2016 | 27.42 | 0.00 | No |
| *BMI* | Melanesia | 2016 | 26.60 | 0.00 | No |
| *BMI* | Micronesia | 2016 | 30.47 | 0.00 | No |
| *BMI* | Northern Africa | 2016 | 26.86 | 0.00 | No |
| *BMI* | Northern America | 2016 | 27.80 | 0.69 | Yes |
| *BMI* | Northern Europe | 2016 | 26.59 | 0.00 | No |
| *BMI* | Polynesia | 2016 | 32.28 | 0.00 | No |
| *BMI* | South-eastern Asia | 2016 | 23.58 | 0.00 | No |
| *BMI* | Southern Asia | 2016 | 23.68 | 0.00 | No |
| *BMI* | Southern Europe | 2016 | 26.62 | 0.00 | No |
| *BMI* | Sub-Saharan Africa | 2016 | 23.59 | 0.00 | No |
| *BMI* | Western Asia | 2016 | 27.80 | 0.64 | Yes |
| *BMI* | Western Europe | 2016 | 25.92 | 0.00 | No |
| *Height* | Australia and New Zealand | 1996 | 171.93 | 0.00 | No |
| *Height* | Central Asia | 1996 | 165.08 | 0.00 | No |
| *Height* | Eastern Asia | 1996 | 166.06 | 0.00 | No |
| *Height* | Eastern Europe | 1996 | 171.45 | 0.00 | No |
| *Height* | Global | 1996 | 166.40 | 0.00 | No |
| *Height* | Latin America and the Caribbean | 1996 | 164.81 | 0.00 | No |
| *Height* | Melanesia | 1996 | 162.36 | 0.00 | No |
| *Height* | Micronesia | 1996 | 161.06 | 0.00 | No |
| *Height* | Northern Africa | 1996 | 164.50 | 0.00 | No |
| *Height* | Northern America | 1996 | 169.06 | 0.00 | No |
| *Height* | Northern Europe | 1996 | 173.22 | 0.00 | No |
| *Height* | Polynesia | 1996 | 169.07 | 0.00 | No |
| *Height* | South-eastern Asia | 1996 | 159.47 | 0.00 | No |
| *Height* | Southern Asia | 1996 | 160.09 | 0.00 | No |
| *Height* | Southern Europe | 1996 | 170.67 | 0.28 | Yes |
| *Height* | Sub-Saharan Africa | 1996 | 162.42 | 0.00 | No |
| *Height* | Western Asia | 1996 | 165.01 | 0.00 | No |
| *Height* | Western Europe | 1996 | 172.50 | 0.00 | No |

**Supplementary Table 6.** Quality metrics of the text mining procedure to assign BED medical conditions. For each BED term, a set of MA samples was copied, and their assigned BED term was manually verified.

| ***BED Term*** | **Total** | **#Correct** | **Accuracy** |
|---|---|---|---|
| *Acne Vulgaris* | 50 | 37 | 74,00 |
| *Acquired Immunodeficiency Syndrome* | 30 | 25 | 83.33 |
| *Acute Lymphocytic Leukemia* | 30 | 16 | 53.33 |
| *Acute Myeloid Leukemia* | 60 | 56 | 93.33 |
| *Adenomatous Polyposis* | 30 | 30 | 100.00 |
| *ADHD* | 60 | 60 | 100.00 |
| *Agitation* | 30 | 0 | 0.00 |
| *Alzheimer Disease* | 30 | 17 | 56.67 |
| *Amyotrophic Lateral Sclerosis* | 30 | 30 | 100.00 |
| *Anaemia* | 30 | 30 | 100.00 |
| *Angina Pectoris* | 30 | 30 | 100.00 |
| *Ankylosing Spondylitis* | 30 | 30 | 100.00 |
| *Anorexia* | 1 | 1 | 100.00 |
| *Anxiety* | 30 | 27 | 90.00 |
| *Arrhythmia* | 30 | 30 | 100.00 |
| *Arthralgia* | 30 | 30 | 100.00 |
| *Asthma* | 30 | 29 | 96.67 |
| *Ataxia* | 30 | 21 | 70.00 |
| *Atherosclerosis* | 30 | 22 | 73.33 |
| *Atrial Fibrillation* | 30 | 30 | 100.00 |
| *Bacterial Infection* | 30 | 30 | 100.00 |
| *Bipolar Disorder* | 30 | 28 | 93.33 |
| *Burning Sensation* | 30 | 0 | 0.00 |
| *Cardiac Valve Disease* | 17 | 4 | 23.53 |
| *Cardiomyopathy* | 30 | 30 | 100.00 |
| *Cholestasis* | 30 | 30 | 100.00 |
| *Chondrosarcoma* | 30 | 30 | 100.00 |
| *Chronic Lymphocytic Leukemia* | 30 | 17 | 56.67 |
| *Chronic Obstructive Pulmonary Disease* | 30 | 23 | 76.67 |
| *Clear Cell Renal Cell Carcinoma* | 30 | 30 | 100.00 |
| *Cognitive Impairment* | 30 | 25 | 83.33 |
| *Colic* | 7 | 1 | 14.29 |
| *Colitis* | 30 | 30 | 100.00 |
| *Colorectal Neoplasms* | 30 | 30 | 100.00 |
| *Conjunctivitis* | 30 | 30 | 100.00 |
| *Constipation* | 30 | 30 | 100.00 |
| *Cough* | 30 | 30 | 100.00 |
| *Crohn Disease* | 30 | 29 | 96.67 |
| *Cystic Fibrosis* | 30 | 30 | 100.00 |
| *Depression* | 30 | 27 | 90.00 |
| *Dermatitis* | 30 | 3 | 10.00 |
| *Dermatitis Atopic* | 30 | 27 | 90.00 |

| | | | |
|---|---|---|---|
| *Diabetes Type I* | 30 | 30 | 100.00 |
| *Diabetes Type Ii* | 60 | 59 | 98.33 |
| *Diabetic Nephropathy* | 30 | 30 | 100.00 |
| *Diabetic Neuropathy* | 6 | 6 | 100.00 |
| *Diabetic Retinopathy* | 18 | 18 | 100.00 |
| *Diarrhoea* | 30 | 30 | 100.00 |
| *DNA Damage Response* | 21 | 17 | 80.95 |
| *Dry Skin* | 1 | 0 | 0.00 |
| *Dyskinesia* | 18 | 10 | 55.56 |
| *Dyspnea* | 30 | 30 | 100.00 |
| *Oedema* | 27 | 27 | 100.00 |
| *Endothelial Dysfunction* | 12 | 0 | 0.00 |
| *Epilepsy* | 30 | 30 | 100.00 |
| *Epistaxis* | 30 | 30 | 100.00 |
| *Erectile Dysfunction* | 12 | 4 | 33.33 |
| *Erythema* | 30 | 13 | 43.33 |
| *Fatigue* | 30 | 26 | 86.67 |
| *Fever* | 30 | 30 | 100.00 |
| *Fibromyalgia* | 30 | 30 | 100.00 |
| *Flushing* | 17 | 0 | 0.00 |
| *Fragile X Syndrome* | 30 | 28 | 93.33 |
| *Gastroesophageal Reflux Disease* | 30 | 29 | 96.67 |
| *Glaucoma* | 30 | 25 | 83.33 |
| *Glioblastoma* | 30 | 29 | 96.67 |
| *Gout* | 2 | 2 | 100.00 |
| *Gynecomastia* | 5 | 5 | 100.00 |
| *Head And Neck Neoplasms* | 30 | 30 | 100.00 |
| *Headache* | 30 | 30 | 100.00 |
| *Heart Failure* | 30 | 28 | 93.33 |
| *Hematuria* | 15 | 14 | 93.33 |
| *Hemochromatosis* | 30 | 25 | 83.33 |
| *Haemorrhage* | 30 | 5 | 16.67 |
| *Hepatitis* | 60 | 60 | 100.00 |
| *Hepatocellular Carcinoma* | 30 | 28 | 93.33 |
| *Hodgkins Lymphoma* | 30 | 0 | 0.00 |
| *Hypercalcemia* | 30 | 10 | 33.33 |
| *Hyperglycemia* | 30 | 0 | 0.00 |
| *Hyperlipidemia* | 30 | 26 | 86.67 |
| *Hypersensitivity* | 30 | 19 | 63.33 |
| *Hypertension* | 30 | 23 | 76.67 |
| *Hyperthermia* | 23 | 23 | 100.00 |
| *Hypertriglyceridemia* | 16 | 16 | 100.00 |
| *Hypocalcemia* | 8 | 6 | 75.00 |
| *Hypocalcemia* | 8 | 0 | 0.00 |

| | | | |
|---|---|---|---|
| *Hypotension* | 7 | 7 | 100.00 |
| *Infection* | 30 | 27 | 90.00 |
| *Inflammation* | 30 | 6 | 20.00 |
| *Leiomyosarcoma* | 30 | 30 | 100.00 |
| *Lipidoses* | 14 | 14 | 100.00 |
| *Liposarcoma* | 30 | 27 | 90.00 |
| *Lower Female Genital Tract Infection* | 19 | 17 | 89.47 |
| *Lung Neoplasms* | 30 | 26 | 86.67 |
| *Lupus Erythematosus Discoid* | 30 | 30 | 100.00 |
| *Lupus Erythematosus Systemic* | 30 | 29 | 96.67 |
| *Lymphopenia* | 7 | 7 | 100.00 |
| *Macular Degeneration* | 30 | 30 | 100.00 |
| *Melanoma* | 30 | 28 | 93.33 |
| *Menopause* | 30 | 23 | 76.67 |
| *Mucositis* | 30 | 30 | 100.00 |
| *Multiple Myeloma* | 30 | 29 | 96.67 |
| *Multiple Sclerosis* | 30 | 30 | 100.00 |
| *Myelodysplastic Syndrome* | 30 | 21 | 70.00 |
| *Myocardial Infarction* | 30 | 18 | 60.00 |
| *Myocarditis* | 5 | 5 | 100.00 |
| *Nasopharyngitis* | 30 | 30 | 100.00 |
| *Nausea Emesis* | 1 | 1 | 100.00 |
| *Necrosis* | 30 | 18 | 60.00 |
| *Nephrosis* | 100 | 87 | 87.00 |
| *Neuroblastoma* | 100 | 100 | 100.00 |
| *Neutropenia* | 9 | 9 | 100.00 |
| *Non-Alcoholic Liver Fatty Disease* | 50 | 50 | 100.00 |
| *Non-Hodgkin Lymphoma* | 50 | 31 | 62.00 |
| *Non-Small Cell Lung* | 50 | 50 | 100.00 |
| *Obesity* | 50 | 44 | 88.00 |
| *Osteoarthritis* | 50 | 29 | 58.00 |
| *Osteoporosis* | 50 | 40 | 80.00 |
| *Osteosarcoma* | 50 | 36 | 72.00 |
| *Ovarian Neoplasms* | 50 | 49 | 98.00 |
| *Pain* | 80 | 68 | 85.00 |
| *Pancreatic Neoplasms* | 49 | 44 | 89.80 |
| *Pancreatitis* | 50 | 50 | 100.00 |
| *Paresthesia* | 3 | 3 | 100.00 |
| *Parkinson Disease* | 50 | 20 | 40.00 |
| *Peptic Ulcer* | 1 | 1 | 100.00 |
| *Periodontitis* | 50 | 50 | 100.00 |
| *Pneumonitis* | 50 | 7 | 14.00 |
| *Primary Myelofibrosis* | 50 | 50 | 100.00 |
| *Prostatic Hyperplasia* | 50 | 50 | 100.00 |

| | | | |
|---|---|---|---|
| *Prostatic Neoplasms* | 50 | 49 | 98.00 |
| *Psoriasis* | 50 | 26 | 52.00 |
| *Pulmonary Arterial Hypertension* | 50 | 37 | 74.00 |
| *Purpura* | 16 | 16 | 100.00 |
| *Renal Cell Carcinoma* | 50 | 27 | 54.00 |
| *Renal Failure* | 50 | 47 | 94.00 |
| *Rheumatoid Arthritis* | 50 | 50 | 100.00 |
| *Rhinitis* | 50 | 50 | 100.00 |
| *Seizures* | 50 | 19 | 38.00 |
| *Septic Shock* | 50 | 40 | 80.00 |
| *Sinusitis* | 1 | 1 | 100.00 |
| *Skin Aging* | 2 | 0 | 0.00 |
| *Skin Pigmentation* | 3 | 3 | 100.00 |
| *Small Cell Lung Neoplasms* | 50 | 15 | 30.00 |
| *Stevens-Johnson Syndrome* | 6 | 6 | 100.00 |
| *Stomach Neoplasms* | 50 | 46 | 92.00 |
| *Stomatitis* | 11 | 11 | 100.00 |
| *Stroke* | 50 | 44 | 88.00 |
| *Tachycardia* | 50 | 0 | 0.00 |
| *Thrombocytopenia* | 50 | 49 | 98.00 |
| *Thrombosis* | 50 | 50 | 100.00 |
| *Thyroid Neoplasms* | 50 | 42 | 84.00 |
| *Tremor* | 35 | 30 | 85.71 |
| *Ulcerative Colitis* | 50 | 46 | 92.00 |
| *Urticaria* | 50 | 5 | 10.00 |
| *Uterine Cervical Neoplasms* | 50 | 39 | 78.00 |
| *Vaginal Bleeding* | 21 | 21 | 100.00 |
| *Vertigo* | 50 | 26 | 52.00 |
| *Waldenstroms Macroglobulinaemia* | 42 | 42 | 100.00 |
| *Weight Gain* | 50 | 46 | 92.00 |
| *Weight Loss* | 50 | 40 | 80.00 |
| *Wilson Disease* | 10 | 10 | 100.00 |

**Supplementary Table 7.** Detailed results of the ML models for the classification of medical conditions according to BED terms. For each BED term, we obtained a set of multiple ML models. The balanced accuracy value reflects the average b. accuracy for that BED term.

| *Family Name* | BED Condition | Number of Models | Balanced Accuracy | CI |
|---|---|---|---|---|
| *Behaviour and Behaviour Mechanisms* | Depression | 40 | 92.56% | [CI 91.29%;93.83%] |
| *Behaviour and Behaviour Mechanisms* | Anxiety | 45 | 93.39% | [CI 92.33%;94.46%] |
| *Cardiovascular Diseases* | Hypertension | 17 | 83.52% | [CI 82.11%;84.94%] |
| *Cardiovascular Diseases* | Myocardial Infarction | 40 | 87.73% | [CI 86.55%;88.91%] |
| *Cardiovascular Diseases* | Stroke | 41 | 90.24% | [CI 89.01%;91.47%] |
| *Cardiovascular Diseases* | Angina Pectoris | 33 | 90.40% | [CI 89.54%;91.27%] |
| *Cardiovascular Diseases* | Atherosclerosis | 35 | 92.97% | [CI 92.12%;93.82%] |
| *Cardiovascular Diseases* | Heart Failure | 42 | 97.13% | [CI 96.26%;98.00%] |
| *Cardiovascular Diseases* | Thrombosis | 31 | 98.47% | [CI 97.73%;99.21%] |
| *Cardiovascular Diseases* | Cardiomyopathy | 43 | 98.92% | [CI 98.46%;99.37%] |
| *Cardiovascular Diseases* | Atrial Fibrillation | 32 | 99.37% | [CI 98.42%;100.32%] |
| *Cardiovascular Diseases* | Hypotension | 40 | 99.38% | [CI 98.66%;100.09%] |
| *Congenital, Hereditary, and Neonatal Diseases and Abnormalities* | Hemochromatosis | 9 | 91.20% | [CI 88.70%;93.71%] |
| *Congenital, Hereditary, and Neonatal Diseases and Abnormalities* | Cystic Fibrosis | 42 | 92.15% | [CI 90.79%;93.50%] |
| *Digestive System Diseases* | Hepatitis | 22 | 85.74% | [CI 84.33%;87.15%] |
| *Digestive System Diseases* | Colitis | 38 | 87.14% | [CI 86.11%;88.18%] |
| *Digestive System Diseases* | Crohn Disease | 37 | 90.06% | [CI 88.86%;91.25%] |
| *Digestive System Diseases* | Mucositis | 28 | 93.29% | [CI 91.66%;94.92%] |
| *Eye Diseases* | Glaucoma | 30 | 92.33% | [CI 91.49%;93.18%] |
| *Eye Diseases* | Macular Degeneration | 46 | 98.91% | [CI 98.44%;99.39%] |
| *Hemic and Lymphatic Diseases* | Primary Myelofibrosis | 23 | 86.16% | [CI 84.69%;87.63%] |
| *Hemic and Lymphatic Diseases* | Thrombocytopenia | 33 | 87.52% | [CI 86.44%;88.61%] |
| *Hemic and Lymphatic Diseases* | Anaemia | 31 | 88.21% | [CI 86.96%;89.46%] |
| *Hemic and Lymphatic Diseases* | Leukopenia | 37 | 95.31% | [CI 94.84%;95.78%] |
| *Infections* | Infection | 9 | 81.57% | [CI 80.66%;82.48%] |
| *Infections* | Bacterial Infection | 43 | 93.68% | [CI 92.53%;94.83%] |
| *Infections* | Pneumonitis | 39 | 97.50% | [CI 96.38%;98.63%] |
| *Mental Disorders* | Bipolar Disorder | 30 | 93.40% | [CI 91.62%;95.18%] |
| *Musculoskeletal Diseases* | Rheumatoid Arthritis | 35 | 87.03% | [CI 85.83%;88.24%] |
| *Musculoskeletal Diseases* | Osteoarthritis | 23 | 90.43% | [CI 88.99%;91.88%] |
| *Musculoskeletal Diseases* | Osteoporosis | 34 | 92.73% | [CI 90.84%;94.62%] |
| *Neoplasms* | Hepatocellular Carcinoma | 21 | 84.29% | [CI 83.27%;85.31%] |
| *Neoplasms* | Neoplasms | 34 | 85.28% | [CI 84.38%;86.17%] |
| *Neoplasms* | Breast Neoplasms | 10 | 85.35% | [CI 82.92%;87.77%] |
| *Neoplasms* | Lung Neoplasms | 18 | 85.36% | [CI 84.04%;86.69%] |
| *Neoplasms* | Osteosarcoma | 25 | 85.83% | [CI 84.66%;87.00%] |
| *Neoplasms* | Renal Cell Carcinoma | 14 | 86.08% | [CI 83.92%;88.23%] |

| | | | | |
|---|---|---|---|---|
| *Neoplasms* | Prostatic Neoplasms | 11 | 86.72% | [CI 84.76%;88.68%] |
| *Neoplasms* | Stomach Neoplasms | 27 | 87.32% | [CI 86.10%;88.53%] |
| *Neoplasms* | Acute Myeloid Leukaemia | 30 | 87.33% | [CI 85.95%;88.70%] |
| *Neoplasms* | Colorectal Neoplasms | 34 | 88.07% | [CI 86.80%;89.34%] |
| *Neoplasms* | Ovarian Neoplasms | 25 | 88.80% | [CI 87.30%;90.29%] |
| *Neoplasms* | Glioblastoma | 29 | 89.70% | [CI 88.48%;90.92%] |
| *Neoplasms* | Head Neck Neoplasms | 42 | 90.51% | [CI 89.21%;91.81%] |
| *Neoplasms* | Melanoma | 35 | 90.60% | [CI 89.55%;91.66%] |
| *Neoplasms* | Liposarcoma | 18 | 91.13% | [CI 89.18%;93.07%] |
| *Neoplasms* | Chondrosarcoma | 30 | 91.38% | [CI 90.29%;92.48%] |
| *Neoplasms* | Multiple Myeloma | 38 | 93.58% | [CI 92.43%;94.72%] |
| *Neoplasms* | Neuroblastoma | 27 | 95.17% | [CI 94.27%;96.07%] |
| *Neoplasms* | Adenomatous Polyposis | 41 | 97.64% | [CI 96.77%;98.50%] |
| *Nervous System Diseases* | Multiple Sclerosis | 40 | 88.91% | [CI 87.60%;90.21%] |
| *Nervous System Diseases* | Alzheimer Disease | 34 | 92.56% | [CI 91.49%;93.63%] |
| *Nervous System Diseases* | Epilepsy | 34 | 93.67% | [CI 92.39%;94.95%] |
| *Nutritional and Metabolic Diseases* | Diabetes Type I | 11 | 84.60% | [CI 82.90%;86.30%] |
| *Nutritional and Metabolic Diseases* | Obesity | 27 | 85.04% | [CI 83.87%;86.20%] |
| *Nutritional and Metabolic Diseases* | Diabetes Type II | 32 | 88.48% | [CI 87.02%;89.93%] |
| *Nutritional and Metabolic Diseases* | Hypercholesterolemia | 36 | 93.65% | [CI 92.49%;94.81%] |
| *Nutritional and Metabolic Diseases* | Hypertriglyceridemia | 13 | 94.36% | [CI 91.98%;96.74%] |
| *Nutritional and Metabolic Diseases* | Hypocalcaemia | 38 | 94.44% | [CI 92.55%;96.34%] |
| *Nutritional and Metabolic Diseases* | Hypercalcemia | 23 | 94.72% | [CI 93.33%;96.11%] |
| *Nutritional and Metabolic Diseases* | Hyperlipidaemia | 21 | 96.03% | [CI 94.05%;98.01%] |
| *Respiratory Tract Diseases* | Dyspnoea | 33 | 99.27% | [CI 98.31%;100.24%] |
| *Signs and Symptoms* | Fever | 24 | 84.09% | [CI 83.02%;85.15%] |
| *Signs and Symptoms* | Pain | 45 | 89.01% | [CI 87.93%;90.10%] |
| *Signs and Symptoms* | Cough | 29 | 89.08% | [CI 87.34%;90.82%] |
| *Signs and Symptoms* | Inflammation | 36 | 89.80% | [CI 88.45%;91.16%] |
| *Signs and Symptoms* | Necrosis | 13 | 90.67% | [CI 88.78%;92.57%] |
| *Signs and Symptoms* | Haemorrhage | 37 | 92.13% | [CI 91.53%;92.72%] |
| *Signs and Symptoms* | Diarrhoea | 38 | 93.49% | [CI 92.33%;94.64%] |
| *Signs and Symptoms* | Fatigue | 38 | 94.14% | [CI 93.33%;94.95%] |
| *Signs and Symptoms* | Constipation | 40 | 94.98% | [CI 93.94%;96.01%] |
| *Signs and Symptoms* | Septic Shock | 48 | 96.05% | [CI 95.23%;96.86%] |
| *Signs and Symptoms* | Ataxia | 31 | 96.24% | [CI 94.50%;97.97%] |
| *Skin and Connective Tissue Diseases* | Erythema | 25 | 92.62% | [CI 91.38%;93.86%] |
| *Skin and Connective Tissue Diseases* | Psoriasis | 42 | 92.79% | [CI 91.63%;93.96%] |
| *Skin and Connective Tissue Diseases* | Dermatitis | 32 | 96.63% | [CI 95.75%;97.51%] |
| *Skin and Connective Tissue Diseases* | Dermatitis Atopic | 6 | 98.97% | [CI 98.00%;99.95%] |

| | | | | |
|---|---|---|---|---|
| *Urogenital Diseases* | Renal Failure | 34 | 87.94% | [CI 86.93%;88.96%] |
| *Urogenital Diseases* | Nephrosis | 26 | 93.46% | [CI 92.08%;94.84%] |
| *Urogenital Diseases* | Haematuria | 38 | 95.98% | [CI 95.58%;96.38%] |
| *Urogenital Diseases* | Diabetic Nephropathy | 35 | 97.00% | [CI 95.77%;98.23%] |

**Supplementary Table 8.** Methods applied for feature selection.

| *Name* | **Description** |
|---|---|
| *Bhattacharyya* | Features are selected with a Bhattacharyya distance-based method, which utilizes a recursive algorithm to obtain the optimal dimension reduction matrix in terms of the minimum upper bound of classification error under a normal distribution. |
| *Elastic Net* | Classification features (variables) are selected using elastic net algorithm with a Lambda parameter (weight of L1 L2 regularization) of 0.1. |
| *Entropy* | The order of feature selection is chosen according to the relative entropy (Kullback-Leibler Distance). |
| *Entropy and correlation of selected features* | The order of feature selection is chosen according to the relative entropy (Kullback-Leibler Distance) and correlation to the previously selected ones. |
| *LASSO* | Classification features (variables) are selected using LASSO (Least Absolute Shrinkage and Selection Operator) L1 regularization regression. |
| *No feature selection applied* | - |
| *Random Forests* | Classification features (variables) are selected using a Random Decision Forest – feature importance. |
| *Random Sets with GLM* | Classification features (variables) are selected using Random Sets using GLM Binomial regression as Classifier. |
| *Relieff* | The Relieff algorithm is used to rank and select the features. It estimates the quality of attributes with and without dependencies among them. It can deal with noisy, incomplete, and multi-class data sets. |
| *Ridge regression feature weights* | Classification features (variables) are selected using Ridge regression |
| *SF linear regression classifier* | Classification features (variables) are selected using the SFFS algorithm. |
| *T-Student test* | Samples without significant differences are excluded. |
| *T-Student test and correlation of features* | Samples without significant differences and that show correlation are excluded. |
| *Wilcoxon rank-sum test* | Samples without significant differences are excluded. |
| *Wilcoxon rank-sum test and features uncorrelation* | Samples without significant differences and that show correlation are excluded. |

**Supplementary Table 9.** Values of trimmed mean and median intensity values of the 40 HK genes used for the background datasets used for spline normalization.

| *Gene* | **Trimmed Mean** | **Median** |
|---|---|---|
| *Q99497* | 10,4864 | 11,2262 |
| *Q15843* | 8,9930 | 9,8123 |
| *P28070* | 9,9129 | 10,5507 |
| *P84090* | 9,6925 | 10,4898 |
| *P28074* | 8,9374 | 9,5148 |
| *P19338* | 9,0749 | 9,6200 |
| *P78344* | 9,2125 | 9,8322 |
| *P84077* | 8,7724 | 9,3304 |
| *P15954* | 10,1681 | 10,9635 |
| *P49773* | 9,8162 | 10,7391 |
| *P50395* | 9,6682 | 10,5082 |
| *P00441* | 10,4296 | 11,3640 |
| *P37108* | 10,4704 | 11,3282 |
| *P61803* | 9,8496 | 10,7300 |
| *P18583* | 8,3646 | 8,8267 |
| *P0CG47* | 11,7611 | 12,8680 |
| *P00558* | 8,9982 | 9,5928 |
| *Q06830* | 10,3853 | 11,1574 |
| *P48556* | 8,6206 | 9,2585 |
| *Q92572* | 8,7227 | 9,2975 |
| *P09429* | 9,2277 | 9,6407 |
| *P25705* | 10,6103 | 11,1946 |
| *Q15046* | 9,1241 | 9,6854 |
| *P00338* | 10,5121 | 11,5263 |
| *P36873* | 9,8293 | 10,8341 |
| *Q15366* | 8,7271 | 9,2709 |
| *P49770* | 7,9992 | 8,4494 |
| *P04844* | 9,3664 | 10,0491 |
| *P0DP23* | 9,3234 | 9,9257 |
| *P14406* | 10,2342 | 11,2600 |
| *P14854* | 9,9354 | 10,8738 |
| *P62875* | 7,9722 | 8,1841 |
| *P0CG48* | 12,0196 | 12,8970 |
| *P10644* | 8,8526 | 9,4748 |
| *O15144* | 9,6915 | 10,5810 |
| *P17844* | 10,5128 | 11,2295 |
| *O95467* | 8,9241 | 9,3446 |
| *P63092* | 8,9207 | 9,3397 |
| *P84996* | 8,9220 | 9,3422 |
| *Q5JWF2* | 8,9227 | 9,3430 |
